# Extracellular DNASE1L3 dysfunction fuels obesity-driven inflammation and metabolic syndrome

**DOI:** 10.64898/2026.03.23.713589

**Authors:** A. Ferriere, A. Roubertie, E. Pisareva, R. Gallo, P. Bandopadhyay, P. Santa, A. Garreau, S. Loizon, D. Brisou, A. Vasilakou, A Cissé, M. Dubois, B. Gatta-Cherifi, P. Zizzari, D. Cota, L. Capuron, N. Castanon, C. Monchaux, J. Izotte, B. Rousseau, L. Mora Charrot, A. Zouine, C. Bianchi, P. Pillet, A. Bibeyran, T. Darde, A. Thierry, N. Djouder, P. Blanco, D. Duluc, D. Ganguly, V. Sisirak

## Abstract

Obesity, a global health crisis affecting 16% of the world population, is characterized by chronic inflammation that contributes to health complications such as type 2 diabetes and metabolic dysfunction-associated steatotic liver disease (MASLD). Emerging evidence suggests that self-DNA released from dying cells aberrantly activates inflammatory responses during obesity. However, the role of extracellular deoxyribonucleases (DNASEs), which at steady state regulate abundance of extracellular self-DNA, remains poorly understood in this context. Here, we show that individuals with obesity exhibit elevated levels of circulating cell-free DNA (cfDNA) with a distinctive end-motif signature, anti-DNASE1L3 autoantibodies and a reduction in circulating DNASE activity. These cfDNA alterations correlate with the severity of obesity and can be corrected by therapeutic intervention such as bariatric surgery. Similarly, mice fed a high-fat diet (HFD) displayed increased cfDNA levels and decreased DNASE activity. Genetic deficiency of the extracellular nuclease DNASE1L3 in mice worsened HFD-induced metabolic complications, including glucose intolerance, insulin resistance, MASLD, and metabolic tissue inflammation. Conversely, targeted supplementation of DNASE1L3 in the liver using adeno-associated viral vectors protected obese mice from developing MASLD and liver inflammation. These findings uncover a novel role of DNASE1L3 in controlling obesity-associated inflammation and its potential therapeutic use for preventing metabolic disease.

## INTRODUCTION

Obesity affects over 16% of the global population and is a major contributor to metabolic syndrome, which is a constellation of conditions including hypertension, dyslipidemia, insulin resistance and abdominal obesity (*1*). Metabolic syndrome significantly increases the risk of chronic diseases such as type 2 diabetes (T2D), cardiovascular disorders, metabolic dysfunction-associated steatotic liver disease (MASLD), cancers and severe COVID-19 (*2–4*). Combined together, these comorbidities collectively account for over 5 million deaths annually and place a substantial burden on quality of life and healthcare systems (*1*).

A defining feature of obesity is chronic, low-grade inflammation of metabolic tissues such as the visceral adipose tissue (VAT) and the liver, a phenomenon termed “metaflammation” (*5*). Metaflammation disrupts systemic metabolic homeostasis and contributes to health complications associated with obesity (*5*). This inflammatory state is driven in part by an accumulation of pro-inflammatory macrophages in the VAT and liver of individuals with obesity (*5*). In the VAT, obesity causes the replacement of anti-inflammatory M2-*like* macrophages by pro-inflammatory M1-*like* macrophages (*6*, *7*), and in the liver, resident macrophages known as Kupffer cells (KC) are progressively replaced by pro-inflammatory monocyte-derived macrophages (*8*, *9*). These inflammatory macrophages secrete cytokines such as tumor necrosis factor (TNF)-α and interleukin (IL)-1β that play a pathogenic role in obesity by promoting insulin resistance, for instance (*5*). These findings are supported by observations demonstrating that the loss of pro-inflammatory cytokines or deletion of pro-inflammatory macrophages ameliorates metabolic complications observed in murine models of diet-induced obesity (DIO) (*10–12*). Yet, anti-cytokine therapies have shown limited success in obese patients (*13*, *14*). Moreover, recent single-cell transcriptomic analyses have revealed a far more complex macrophage landscape within the VAT and liver than previously appreciated, emphasizing the need to better understand the regulatory mechanisms underlying metaflammation in obesity (*15*, *16*).

Cellular stress induced by obesity leads to the release of damage-associated molecular patterns (DAMPs) from the cells of the metabolic tissues, which activate pattern recognition receptors (PRRs) and contribute to metaflammation (*17–19*). Among these DAMPs, emerging evidence highlights a prominent role for endogenous cell-free DNA (cfDNA), which accumulates during obesity (*20*, *21*). This accumulation results from increased adipocyte and hepatocyte death (*22*, *23*), as well as enhanced neutrophil extracellular trap (NET) formation through the process of NETosis (*24*). This cfDNA, either associated with microparticles (MPs) derived from dying cells (*23*) or bound to proteins such as HMGB1 (*25*), activates DNA-sensing receptors including Toll-like receptor 9 (TLR9) and cyclic GMP-AMP synthase (cGAS), leading to the production of inflammatory cytokines and type I interferons (IFN-I) and ultimately to the metaflammation caused by obesity. Supporting these findings, mice lacking TLR9 or cGAS and their signaling pathways are protected from obesity-induced inflammation and metabolic complications (*22–27*). These responses are primarily mediated by innate immune cells within metabolic tissues, including plasmacytoid dendritic cells (pDCs) (*25*, *28*), conventional type I dendritic cells (cDC1) (*29*), and macrophages (*23*, *27*), all of which play key roles in metaflammation-induced by obesity (*20*).

Despite the central role of cfDNA in metaflammation, the mechanisms regulating its levels and immunostimulatory potential during obesity remain poorly understood. Circulating deoxyribonucleases (DNASEs), particularly DNASE1 and DNASE1-*Like* 3 (DNASE1L3), are key enzymes that clear extracellular cfDNA and prevent its ability to cause inappropriate immune activation (*30*). While DNASE1 digests “naked” DNA, DNASE1L3 preferentially degrades membrane-bound and nucleosome-associated DNA (*31–33*). Importantly, DNASE1L3 deficiency, but not DNASE1, leads to systemic autoimmunity in form of lupus both in mice and humans due to the accumulation and persistence of immunostimulatory DNA complexed with MPs (*33–35*). Intriguingly, humans and mice prone to systemic autoimmunity and cfDNA accumulation were reported to be more susceptible to metabolic syndromes induced by obesity (*36*, *37*). Furthermore, studies have shown that total circulating DNASE activity is reduced in murine models of obesity (*24*) and hypercholesterolemia (*38*, *39*), potentially impairing DNA clearance. However, DNASE1 supplementation in obese mice did not show any therapeutic effects, possibly due to its restricted substrate specificity or functional inhibition by obesity-related factors (*24*).

In this study, we investigated how DNASE1L3 modulates cfDNA accumulation and metaflammation in obesity. We found that obese patients display elevated and altered profile of nuclear cfDNA, reduced DNASE activity, and the presence of DNASE1L3-specific auto-antibodies. Notably cfDNA accumulation correlated with disease severity and was reduced following bariatric surgery. In mouse models of DIO, *Dnase1l3-*deficiency exacerbated weight gain, heightened adipose and liver inflammation (metaflammation), and worsened MASLD. Conversely, liver-targeted adeno-associated virus (AAV) delivery of human DNASE1L3 restored circulating DNASE activity and attenuated MASLD progression by limiting inflammation induced by HFD. These findings identify DNASE1L3 as a critical regulator of cfDNA homeostasis and inflammation in obesity and highlight its therapeutic potential for metabolic and hepatic complications.

## RESULTS

### Obese patients display elevated levels of circulating cfDNA with a distinctive signature

Obesity has been shown to induce adipocyte and hepatocyte death, leading to the release of their genetic material into circulation (*22*, *23*). Based on this, we investigated whether obesity alters both the quantity and quality of circulating cfDNA. Our study included 42 obese patients (OB) and 43 age- and sex-matched healthy donors (HD), detailed in **Table 1**. Using a qPCR assay targeting ALU repetitive elements (*40*), we found that OB patients displayed significantly higher levels of nuclear (n) cfDNA compared to HD (49.46 ng/ml vs. 21.85 ng/ml, p = 0.0004) (**Fig.1A**). In contrast, mitochondrial (mt) cfDNA levels remained unchanged between the two groups, as measured by a qPCR assay specific for mitochondrial genes (**Fig.1B**). Consequently, OB patients exhibited a reduced mitochondrial-to-nuclear cfDNA ratio compared to HD (**Fig. S1A**). We also observed that the excess nuclear cfDNA in OB patients was primarily found in the soluble plasma fraction rather than in MPs shed by dying cells, as purified by high-speed centrifugation (**Fig.1C**). Interestingly, OB patients had significantly fewer circulating MPs than healthy donors (**Fig. S1B**). In fact, 72% of nuclear cfDNA in OB patients was found in the soluble fraction, while only 28% was MP-associated (**Fig. S1C**). In contrast, cfDNA was evenly distributed across both fractions in HD (**Fig. S1C**). Despite the reduced number of MPs, the nuclear DNA content within these MPs was significantly higher in OB patients than in HD subjects (0.03 pg/ml vs. 0.008 pg/ml, p = 0.034) (**Fig.1D**). This suggests that both soluble and MP-associated cfDNA levels are elevated during obesity. Since NETs have been proposed as additional source of cfDNA in obesity (*41*), we assessed circulating levels of myeloperoxidase (MPO) and neutrophil elastase (NE). We found no overall increase in MPO or NE in OB individuals (**Fig. S1D**), suggesting no widespread neutrophil activation or death. However, unlike in HD, nuclear cfDNA levels in OB patients correlated significantly with MPO and NE, implying a neutrophil contribution to cfDNA accumulation (**Fig. S1E**). Next, we explored links between nuclear cfDNA and obesity-related metabolic complications. Nuclear cfDNA levels positively correlated with the body mass index (BMI), insulin levels, and insulin resistance (HOMA-IR) (**Fig.1E**), but not with LDL, triglycerides (TG), alanine aminotransferase (ALAT), or glycemia (**Fig. S1F**). In a longitudinal cohort of 25 OB patients undergoing bariatric surgery (**Table 2**), we found that surgery-induced weight loss significantly reduced nuclear cfDNA levels (**Fig.1F**), reinforcing the association between cfDNA accumulation and obesity severity. Finally, we analyzed the quality of cfDNA in OB individuals compared to HD and patients carrying a null mutation in *DNASE1L3* (D1L3^null^) by sequencing. Obese patients and HD displayed a similar overall size distribution, distinct from D1L3^null^ patients, with 166 bp mono-nucleosomal fragments predominating (**Fig.1G**). However, OB individuals showed a higher frequency of large cfDNA fragments (>250pb) than HD, reaching levels comparable to D1L3^null^ patients (**Fig.1G-H**), indicating an impaired cfDNA fragmentation as it was previously reported in D1L3^null^ individuals (*42–44*). Because cfDNA end-motif are regulated by circulating DNASEs (*42–44*) we next examined their specific features. Principal component analysis (PCA) revealed a distinct clustering of 5’ end-motifs (4bp) between OB patients, HD and D1L3^null^ patients (**Fig.1I**). While the specific end-motifs depletion characteristic of D1L3^null^ patients was not observed in obesity (**Fig. S1G**), OB patients showed reduced GC content of cfDNA end-motifs, intermediate between HD and D1L3^null^ patients (**Fig 1J**). Together these finding demonstrate that obesity leads to the accumulation of nuclear cfDNA in the circulation with a distinct fragmentation and end-motif signature, likely reflecting impaired DNASE activity. This nuclear cfDNA buildup may contribute to obesity-associated inflammation given its association with obesity severity.

**Figure 1.**
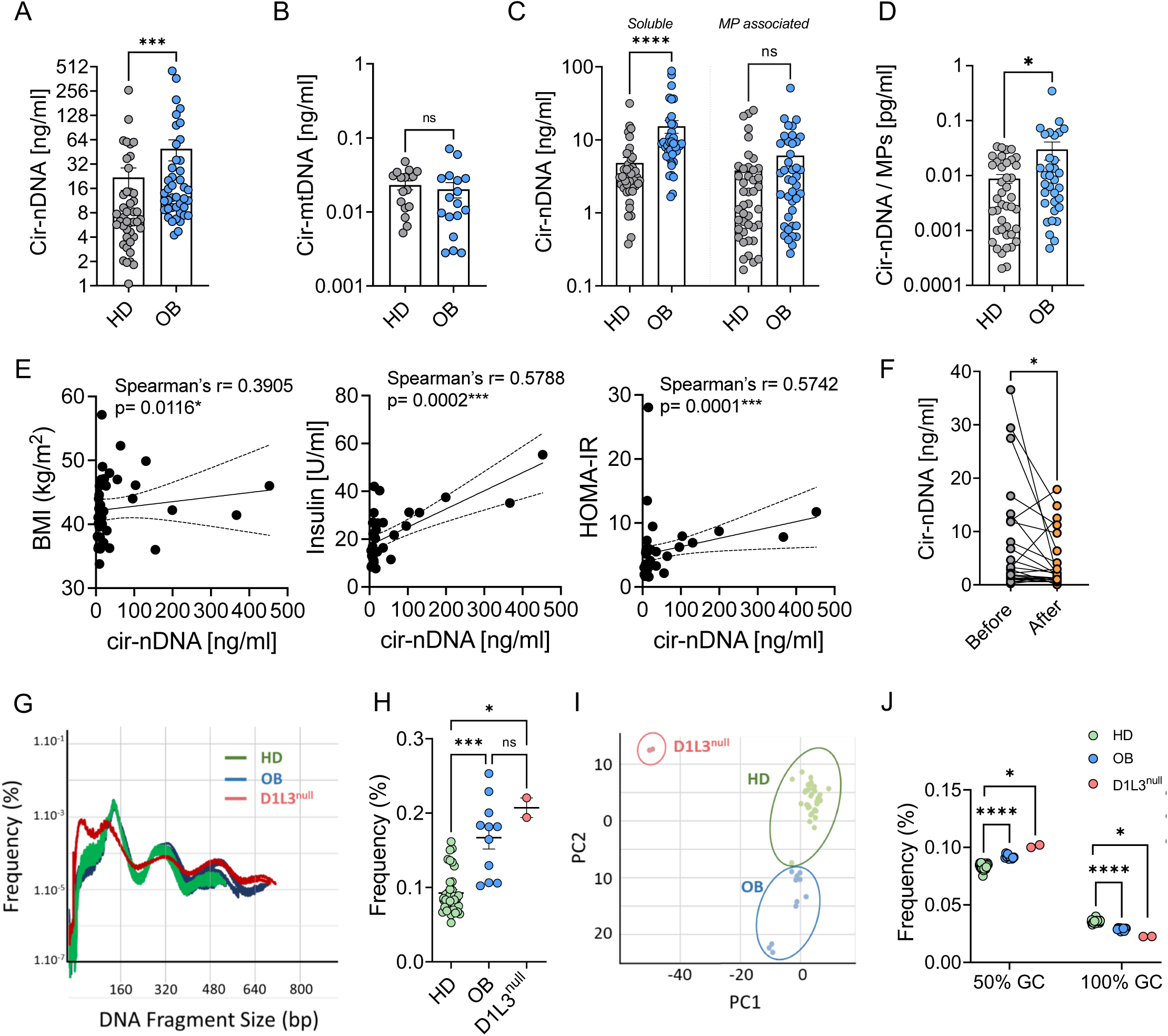
Obese patients display elevated levels of circulating cfDNA with a distinctive signature. (**A-B**) Circulating levels of nuclear (n) DNA (A) and mitochondrial (mt) DNA (B) in the plasma from obese patients (OB) and healthy donors (HD). (**C**) Circulating nDNA levels in soluble (MP⁻) and microparticle-associated (MP⁺) fractions of the plasma from OB and HD. (**D**) Ratio of circulating nDNA content per microparticle (MP⁺) in plasma from OB and HD. (**E**) Correlation of circulating nDNA levels with body mass index (BMI), insulin levels, and HOMA-IR, analyzed by linear regression. Spearman’s r and p-values are indicated on each graph. (**F**) Circulating nDNA levels in obese patients before and 3 to 12 months after bariatric surgery. (**G-H**) Overall size distribution of cfDNA (**G**) and frequency of circulating DNA fragments >250bp (**H**) from HD (n=48), OB (n=11), and DNASE1L3-null (D1L3^null^) individuals (n=2), as determined by sequencing. (**I**) PCA of 5′ end-motif signatures in cfDNA from HD, OB, and DNASE1L3-null individuals, derived from sequencing. (**J**) Proportion of GC-enriched 4bp end motifs in HD (n=37), OB (n=11), and D1L3^null^ individuals (n=2), based on sequencing data. Data are presented as mean ± SEM. Statistical significance was determined using unpaired two-tailed t-tests or one way ANOVA. Significance is denoted as follows: ns = not significant; *p ≤ 0.05; ***p ≤ 0.001; ****p ≤ 0.0001.

**Table 1.** Characteristics of the cohort 1 of obese patients and healthy donors.

| Variables | Patients (n=42) |  | Controls (n=43) |  | p-value |
| --- | --- | --- | --- | --- | --- |
|  | N (%) / x (SD) | Range | N (%) / x (SD) | Range |  |
| **Age** | 41.71 (10.99) | 22.0 - 65.0 | 37.46 (23.26) | 18.0 - 61.0 | 0,14 |
| **Gender** |  |  |  |  |  |
| Female | 33 (78.6) | / | 32 (74.4) | / | / |
| Male | 9 (21.4) | / | 11 (25.6) | / | / |
| **BMI** | 42.47 (5) | 33.8 - 57.13 | 23.26 (3.07) | 17.8 - 29.3 | <0,0001 |
| **BMI max** | 44.64 (5.67) | 37 - 64.0 | / | / | / |
| **Glycemia (g/l)** | 1.04 (0.74) | 0.76 - 5.6 | / | / | / |
| **Insulinemia (μU/ml)** | 21.33 (11.28) | 7.7 - 55.3 | / | / | / |
| **HOMA-IR** | 5.61 (0.8) | 1.6 - 28.2 | / | / | / |
| **ALAT (U/l)** | 30.05 (17.11) | 8 - 93 | / | / | / |
| **TG (g/l)** | 1.17 (0.52) | 0.39 - 2.45 | / | / | / |
| **LDL (g/l)** | 1.3 (0.40) | 0.51 - 2.7 | / | / | / |

**Table 2.** Characteristics of the cohort of obese patients that underwent bariatric surgery.

| Variables | Patients (n=25) |  | p-value |
| --- | --- | --- | --- |
|  | N (%) / x (SD) | Range |  |
| **BMI before surgery (kg/m<sup>2</sup>)** | 41.55 (5.27) | 35.2 - 59.1 | / |
| **BMI after surgery (kg/m<sup>2</sup>)** | 30.28 (5.09) | 22.84 - 43.36 | / |
| **Weight loss (%)** | 27.19 (7.71) | 13.83 - 40 | <0,0001 |
| **Time to follow-up (months)** | 7.2 (3.2) | 3.0 - 12.0 | / |

### Obese patients display a reduced circulating DNASE activity and anti-DNASE1L3 autoantibodies

Circulating cfDNA levels and their molecular characteristics are tightly regulated by extracellular/circulating DNASEs (*42*). In particular, DNASE1L3, which is produced predominantly by innate immune cells (*33*) and liver endothelial cells (*45*), has been shown to control both cfDNA fragment size and end-motif signatures (*42–44*). Our cfDNA analysis revealed that OB individuals accumulate cfDNA with distinct molecular features, suggestive of impaired DNASE1L3 regulation. To explore this, we first assessed DNASE1L3 expression in peripheral blood mononuclear cells (PBMCs) of OB individuals. Due to the lack of specific antibodies for DNASE1L3, we used a Flow-FISH approach to quantify its mRNA expression. We found that total DNASE1L3 expression in PBMCs was unchanged in OB patients compared to HD (**Fig.2A**). As expected, DNASE1L3 expression was confined primarily to pDCs and, to a lesser extent, monocytes (**Fig.S2A**). Obesity did not affect DNASE1L3 expression in any of the immune cell subsets we analyzed (**Fig.S2B**). We then evaluated DNASE1L3 expression in visceral adipose tissue (VAT) from OB (n=20) and lean (n=10) individuals. Interestingly, DNASE1L3 expression was significantly higher in the VAT of OB patients (**Fig.2B**), likely reflecting immune cell infiltration induced by obesity (*46*). Further profiling of VAT-infiltrating immune cells revealed that, while DNASE1L3 expression was reduced in pDCs compared to their blood counterparts, it was elevated in conventional DC and monocytes within the VAT (**Fig.S2C**). This analysis could not be extended to HD due to the limited immune cell yield from lean VAT. Additionally, we differentiated monocyte-derived macrophages *in vitro* under M-CSF and IL-4 to model anti-inflammatory M2-*like* macrophages. DNASE1L3 expression was high in M2-*like* macrophages but was markedly suppressed by IFN-I, which promotes the pro-inflammatory M1-*like* phenotype (*25*) (**Fig.S2D**). These findings suggest that DNASE1L3 expression is increased in the VAT and that its distribution shifts across immune cell types compared with blood. Beyond expression analysis, we assessed systemic DNASE enzymatic activity in plasma from OB patients and healthy donors using a single radial enzyme-diffusion (SRED) assay. Although limited to a small number of patients (n=3), the SRED assay revealed reduced DNASE activity in OB individuals (**Fig.2C**). To validate this observation in our whole cohort, we developed an alternative assay using QuantiT-PicoGreen™ assay to quantify dsDNA degradation by the plasma. This assay confirmed a significant reduction in global DNASE activity in OB patients compared to HD (**Fig.2D**). Moreover, DNASE activity negatively correlated with circulating nuclear cfDNA levels in OB patients (**Fig.2E**), but showed no significant associations with other metabolic markers such as LDL, TG, ALAT, insulin, or glucose (**Fig.S2E**). Furthermore, we observed that bariatric surgery did not impact the overall circulating DNASE activity (**Fig.S2F).** Given this evidence of reduced DNASE function in obesity, we next asked whether autoantibodies might be involved, as they are known to inhibit DNASE enzymes in autoimmune settings (*47*, *48*). Since obesity is linked to heightened B cell autoreactivity (*24*), we quantified circulating antibodies against DNASE1 and DNASE1L3, which are the nucleases responsible of the totality of DNASE activity detected in the circulation (*49*). While anti-DNASE1 antibodies were undetectable in obese individuals (**Fig.2F**), we observed a significant increase in anti-DNASE1L3 autoantibodies in OB patients compared to HD (**Fig.2G**). Specifically, 15 out of 59 obese individuals tested positive for these autoantibodies. However, the weight loss induced by bariatric surgery did not affect anti-DNASE1L3 autoantibodies (**Fig. S2G**). We confirmed the presence of these autoantibodies in an independent cohort detailed in **Table 3** comprising 25 OB individuals and 10 HD (**Fig. 2H**). In this second cohort, more than half of OB patients were positive for anti-DNASE1L3 autoantibodies, and their levels positively correlated with BMI (**Fig. 2I**). Altogether, these findings suggest that obesity impairs DNASE1L3 activity, likely through inhibitory autoantibodies, ultimately contributing to the abnormal accumulation and altered profile of cfDNA in OB individuals.

**Figure 2.**
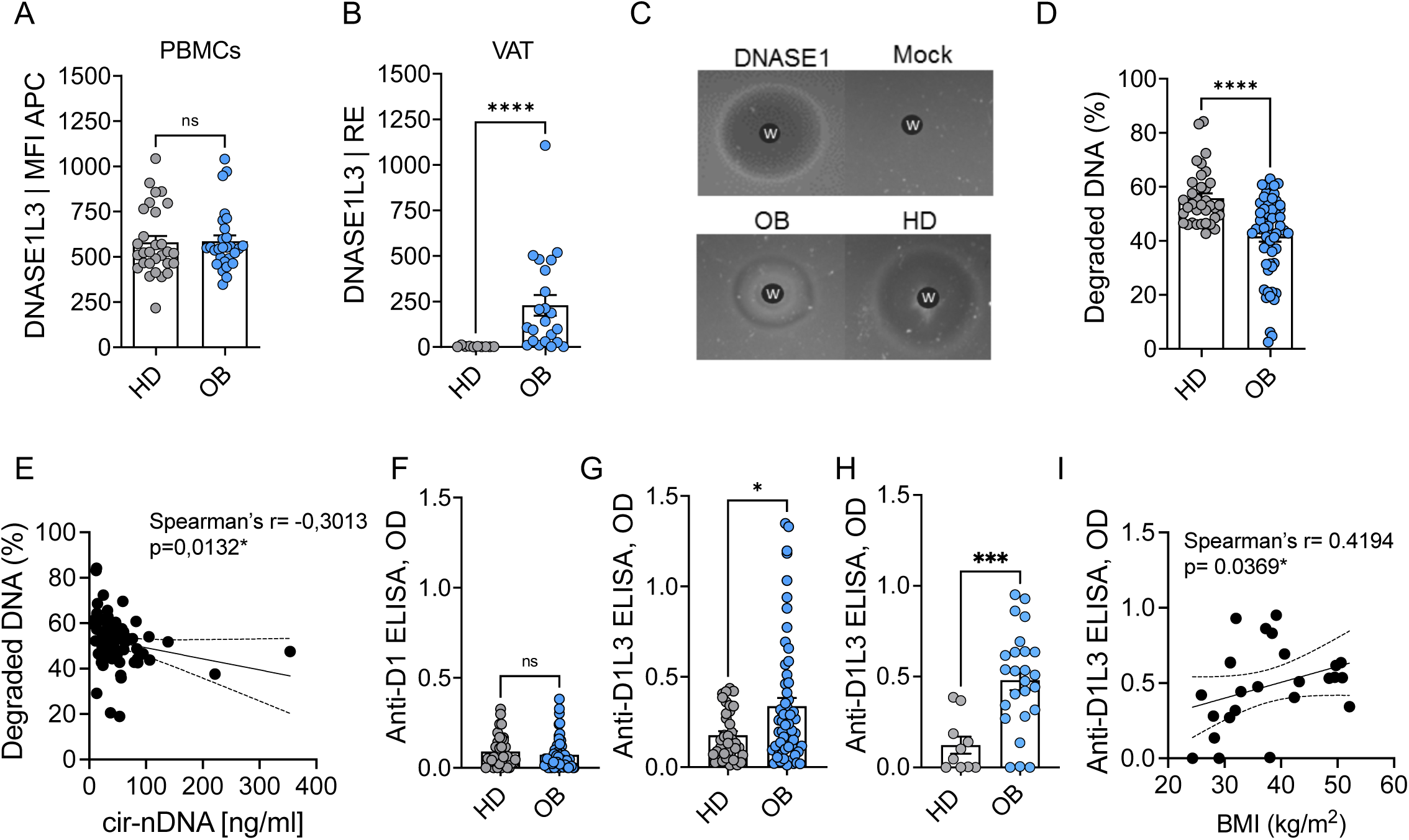
Obese patients exhibit reduced circulating DNASE activity. (**A**) Mean fluorescence intensity (MFI) of DNASE1L3 mRNA in peripheral blood mononuclear cells (PBMCs) from obese patients (OB) and healthy donors (HD), measured by Flow-FISH. (**B**) DNASE1L3 mRNA expression in visceral adipose tissue (VAT) from OB and HD, assessed by quantitative PCR. (**C**) Total circulating DNASE activity in OB and HD, quantified using the single radial enzyme-diffusion (SRED) assay. (**D**) Percentage of degraded naked DNA following incubation with plasma from OB or HD. (**E**) Correlation between plasma DNASE activity (as % degraded DNA) and circulating nuclear (n) cfDNA levels in obese individuals. r and p-values determined by Spearman’s test are shown for each correlation. (**F**) Plasma levels of anti-DNASE1 autoantibodies in OB and HD, measured by ELISA. (**G-H**) Plasma levels of anti-DNASE1L3 autoantibodies in OB and HD in two independent cohorts, measured by ELISA. (**I**) Correlation between plasma anti-DNASE1L3 antibody levels and the BMI in obese individuals. r and p-values determined by Spearman’s test are shown for each correlation. Data are presented as mean ± SEM. Statistical analysis was performed using unpaired two-tailed t-tests. Significance is denoted as follows: ns = not significant; *p ≤ 0.05; ****p ≤ 0.0001.

**Table 3.**
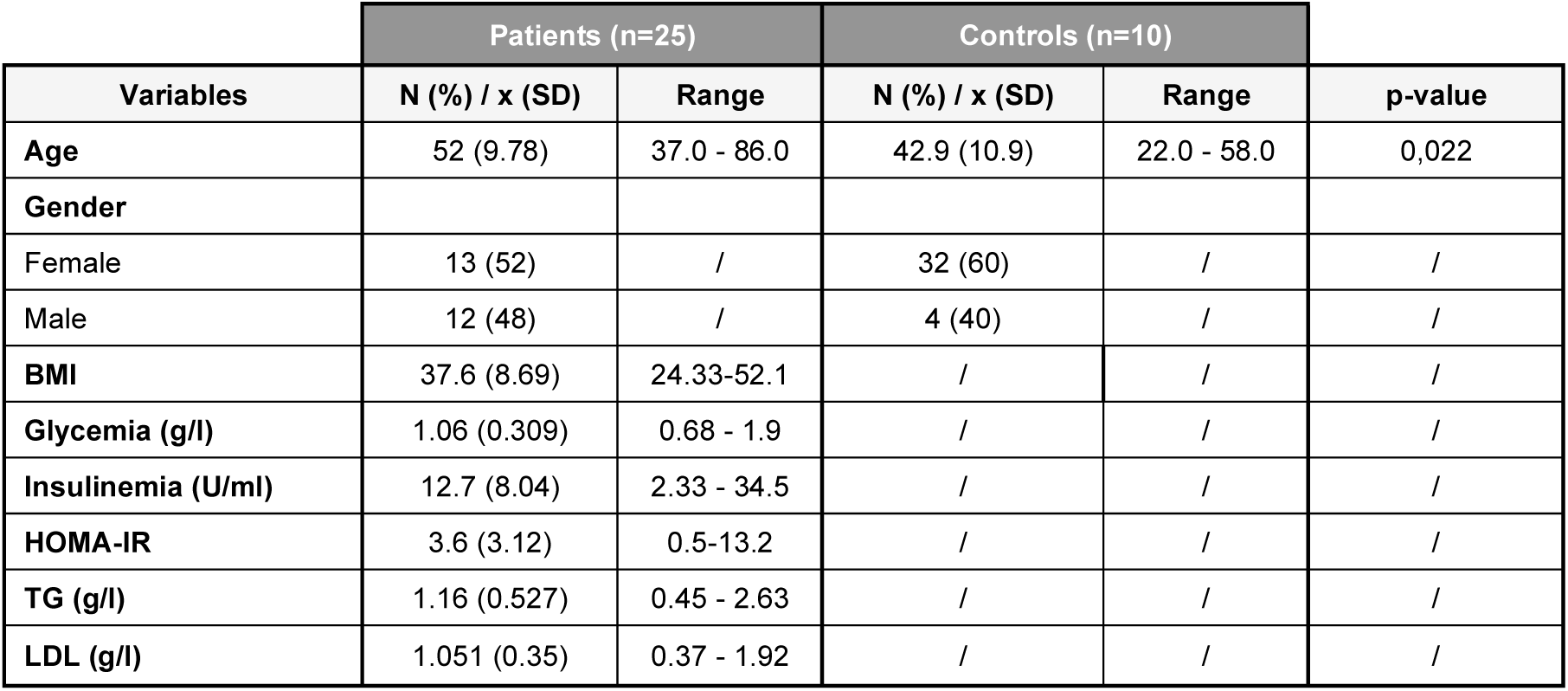
Characteristics of the cohort 2 of obese patients and healthy donors.

### The deficiency in DNASE1L3 exacerbates weight gain in a mouse model of diet-induced obesity

We next investigated whether the alterations observed in OB individuals could be reproduced in a mouse model of DIO. To this end, wild-type (WT) mice were fed for 12 weeks either a normal diet (ND) or a HFD containing 45% of fat, which closely reflects the macronutrient composition and chronic metabolic stress of a Western diet (*50*). We then assessed circulating cfDNA levels and DNASE activity. Mice on HFD exhibited reduced circulating DNASE activity, as evidenced by the diminished ability of their plasma to degrade DNA (**Fig. 3A**). In parallel, HFD-fed mice showed elevated levels of circulating cfDNA of nuclear origin (**Fig. 3B**), whereas mitochondrial cfDNA levels remained unchanged (**Fig. 3C**). Given that DNASE1L3 is a major DNASE regulating the quantity and the quality of circulating cfDNA (*42–44*) and that a subset of OB patients develops anti-DNASE1L3 antibodies, we next examined the impact of *Dnase1l3*-deficiency in a mouse model of DIO. WT and *Dnase1l3*-deficient (KO) male mice were fed either ND or HFD containing 45% fat. Cumulative food intake, measured over 7 days, was similar between WT and *Dnase1l3*-KO mice (**Fig. S3A**). Under ND conditions, *Dnase1l3*-KO mice displayed a 24-hour food intake comparable to WT mice (**Fig. S3B**). Interestingly, despite similar food intake, *Dnase1l3*-KO mice exhibited an increased respiratory exchange ratio (RER) during the light phase (**Fig. S3G**), indicating altered fasting-associated substrate utilization under physiological conditions. In contrast, under HFD conditions, *Dnase1l3*-KO mice showed a redistribution of feeding behavior, characterized by increased food consumption during the light phase and reduced intake during the dark phase (**Fig. S3C**), without changes in total 24-hour intake, indicating a diet-dependent alteration in feeding rhythms rather than caloric intake per se. This shift in feeding behavior in HFD-fed *Dnase1l3*-KO mice was accompanied by a reduction in energy expenditure during the late dark phase (**Fig. S3D**). However, 24-hour oxygen consumption (**Fig. S3E**), carbon dioxide production (**Fig. S3F**), RER (**Fig. S3G**), and energy expenditure (**Fig. S3H**) of *Dnase1l3*-KO mice remained comparable to WT controls, consistent with HFD-induced metabolic inflexibility masking genotype-dependent differences in substrate utilization. Despite these relatively modest alterations in metabolic cage parameters, *Dnase1l3-*KO mice gained significantly more weight than WT mice when fed a HFD (**Fig. 3D**). This difference became statistically significant after 6 weeks of HFD and persisted throughout the feeding period (**Fig. 3E**). After 12 weeks, *Dnase1l3*-KO mice had gained approximately 15% more weight than their WT counterparts (12.4 ± 3.6 g *vs* 8.1 ± 3.1 g) (**Fig. 3F**). In contrast, no significant differences in body weight were observed between WT and KO mice fed a ND (**Fig. 3D–F**). HFD-fed *Dnase1l3*-KO mice also exhibited elevated epididymal (visceral-EAT) and inguinal (subcutaneous-IAT) adipose tissue depots compared to WT controls (**Fig. 3G–H**). This was accompanied by larger epididymal adipocyte perimeter and surface area (**Fig. 3I–K**). Body composition analysis confirmed an increase in total fat mass (16,6 g ± 3,09 *vs* 10,9 g ± 4,43, p= 0.0040) and a reduction in total lean mass (19,14 g ± 1,44 *vs* 22,14 ± 2,46 p=0.0007) in *Dnase1l3*-KO mice compared to WT mice fed a HFD (**Fig. 3L–M**). Similar findings were obtained in male mice fed a 60% HFD (**Fig. S3I–M**), as well as in female mice fed a 60% HFD (**Fig. S3N–R**). Together, these results demonstrate that DNASE activity is reduced in mouse models of DIO, and that *Dnase1l3* deficiency exacerbates weight gain and adiposity in response to HFD, independent of sex.

**Figure 3.**
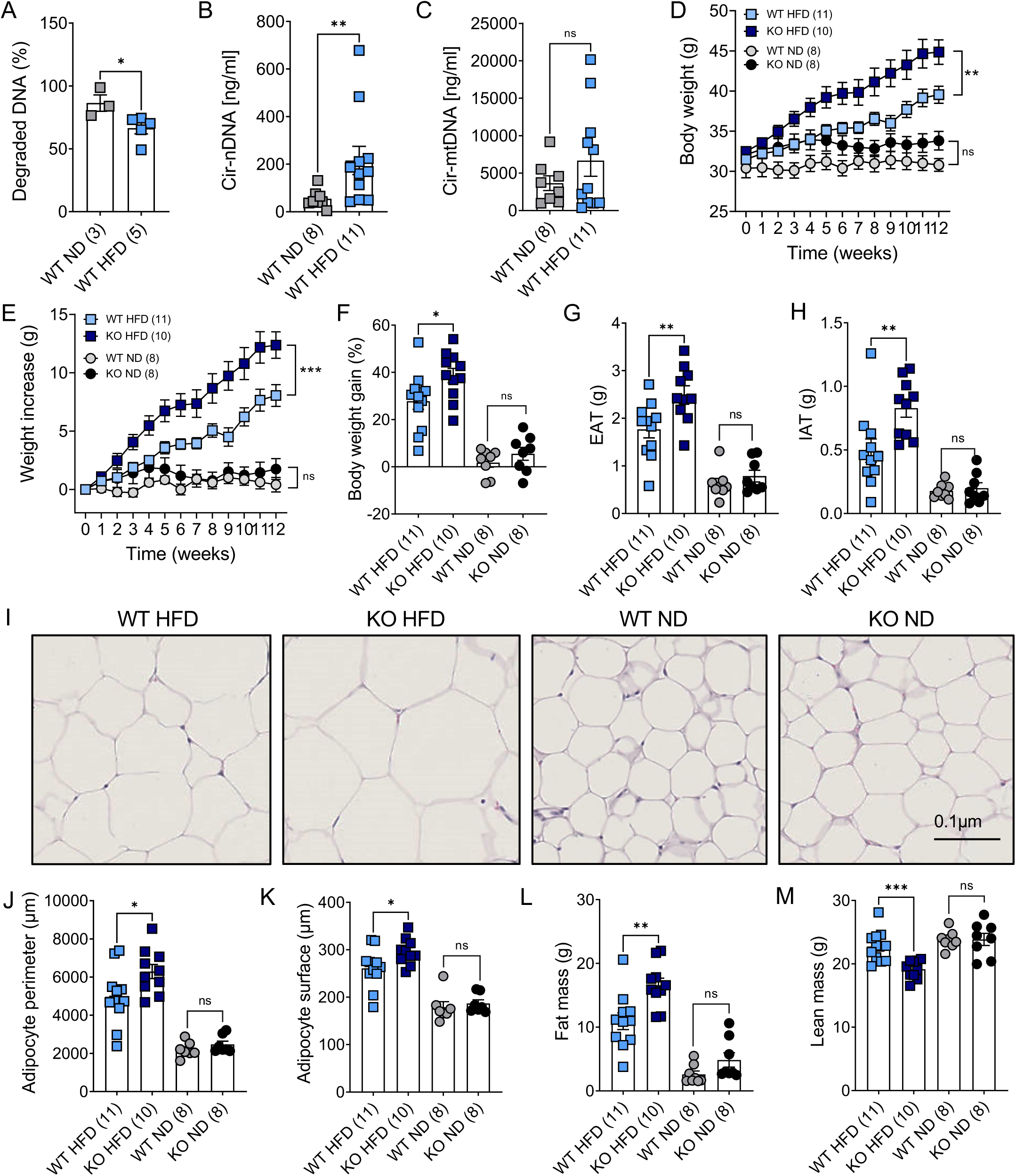
*Dnase1l3-*deficiency exacerbates weight gain in male mice fed a 45% HFD. **(A)** Percentage of degraded naked DNA after incubation with plasma from WT mice fed a normal diet (ND) or 45% HFD for 12 weeks. (**B-C**) Circulating nuclear (**B**) and mitochondrial (**C**) DNA levels in the plasma of WT mice fed ND or 45% HFD for 12 weeks. (**D**) Weekly body weight measurements of WT and *Dnase1l3-*KO mice fed a ND or 45% HFD. (**E**) Weight gain of WT and *Dnase1l3*-KO mice fed a ND or 45% HFD. (**F**) Cumulative body weight gain in WT and *Dnase1l3*-KO mice after 12 weeks on ND or 45% HFD. (**G–H**) Mass of the epididymal adipose tissue (EAT) (**G**) and inguinal adipose tissue (IAT) (**H**) in WT and *Dnase1l3*-KO mice after 12 weeks on ND or 45% HFD. (**I**) Representative H&E-stained images of EAT from the indicated genotypes and diets. Scale bar = 0.1 μm. (**J-K**) Quantification of adipocyte perimeter (**J**) and surface (**K**) from 30 randomly selected cells per genotype and diet. (**L-M**) Fat (**L**) and lean (**M**) mass of WT and *Dnase1l3*-KO mice as measured by MRI after 12 weeks on ND or 45% HFD. Data were pooled from two independent experiments with the number of mice indicated. Results are presented as mean ± SEM. Statistical analyses were performed using unpaired two-tailed t-tests and two-way ANOVA. Significance is denoted as: ns = not significant; *p ≤ 0.05; **p ≤ 0.01; ***p ≤ 0.001.

### The deficiency in DNASE1L3 exacerbates metabolic syndrome and liver disease induced by obesity

Obesity leads to multiple health complications that are recapitulated in murine models of DIO, including glucose intolerance, insulin resistance, and MASLD. In our experimental setting, male mice fed a 45% HFD for 12 weeks developed greater glucose intolerance compared to those on ND (**Fig. 4A–B**). Fasting blood glucose levels were significantly higher in *Dnase1l3*-KO mice than in WT controls fed a 45% HFD (2.24 ± 0.4 *vs* 1.77 ± 0.4 mmol/l, p = 0.0096) (**Fig. 4C**). In addition, *Dnase1l3*-KO mice displayed elevated fasting insulin levels and an increased insulin resistance index, as measured by the HOMA-IR score, compared to WT mice under the same dietary conditions (**Fig. 4D–E**). Despite these metabolic alterations, blood lipid profiles were not significantly affected by the 45% HFD. Both WT and *Dnase1l3*-KO mice exhibited comparable levels of circulating LDL, total cholesterol, and TG regardless of diet (**Fig. 4F**). Similar observations were made in male mice fed a 60% HFD. Interestingly, *Dnase1l3*-deficiency exacerbated glucose intolerance (**Fig. S4A–C**), insulin resistance (**Fig. S4D–G**) in these mice after 6 weeks on the 60% HFD. However, these differences were no longer statistically significant after 12 weeks of 60% HFD feeding (**Fig. S4I–O**). Because the 60% HFD contains a higher fat content than the 45% HFD, it led to elevated circulating lipid levels, particularly in *Dnase1l3*-KO mice (**Fig. S4H-P**). These results suggest that *Dnase1l3*-deficiency accelerates the onset of metabolic dysfunction induced by HFD in male mice. In contrast, female mice remained metabolically protected despite 60% HFD exposure. Although *Dnase1l3*-KO females gained more weight than WT females on HFD (**Fig. S3N-O**), they did not exhibit significant changes in glucose metabolism or insulin sensitivity (**Fig. S4S-W)**, indicating resistance to HFD-induced metabolic dysfunction, as previously described (*50*). We next assessed whether a deficiency in DNASE1L3 affected liver pathology associated with obesity. While liver mass was not significantly affected by 45% HFD feeding (**Fig. 4G**), circulating ALAT levels, a marker of liver damage, were significantly elevated in *Dnase1l3*-KO mice compared to WT mice after both 6 and 12 weeks of HFD (**Fig. 4H-I**). Histological analysis further revealed that hepatic steatosis developed only in *Dnase1l3*-KO mice under these conditions (**Fig. 4J-K**). Mice fed a 60% HFD developed more pronounced liver pathology than those on the 45% HFD, with significantly greater steatosis observed in *Dnase1l3*-deficient animals (**Fig. S4Q-R**). Similarly, *Dnase1l3*-deficiency also aggravated liver steatosis in female mice fed a 60% HFD (**Fig. S4X**). Taken together, these results indicate that *Dnase1l3*-deficiency exacerbates both metabolic complications and liver pathology associated with obesity, in multiple models of DIO.

**Figure 4.**
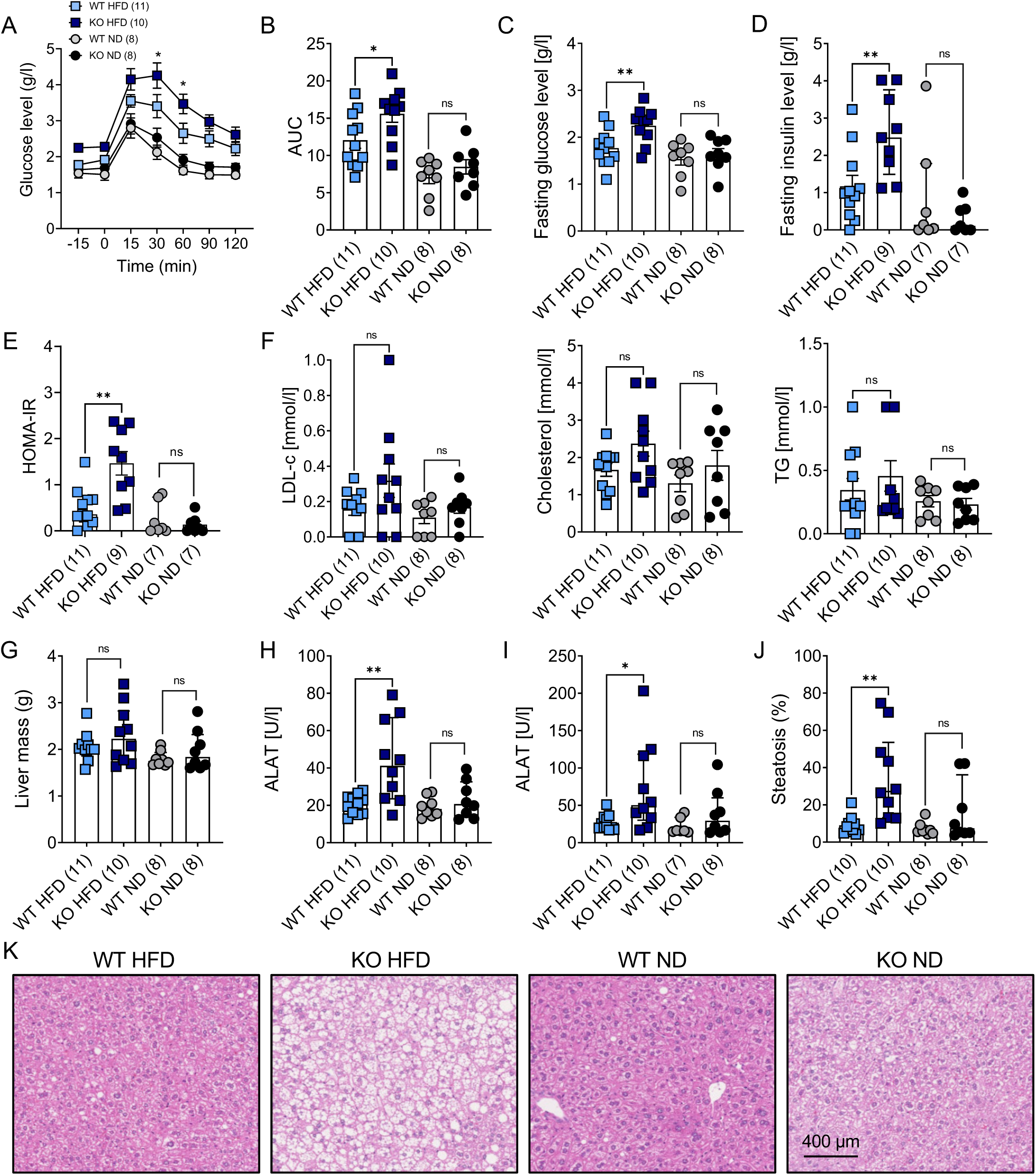
*Dnase1l3*-deficiency exacerbates metabolic syndrome and MASLD in HFD (45%) fed male mice. **(A)** Glucose tolerance test (GTT) in WT and *Dnase1l3*-KO mice after 12 weeks on a normal diet (ND) or 45% HFD. (**B**) Area under the curve (AUC) analysis of the GTT in WT and *Dnase1l3*-KO mice. (**C–D**) Fasting blood glucose (**C**) and insulin levels (**D**) in WT and *Dnase1l3*-KO mice after 12 weeks on ND or 45% HFD. (**E**) Homeostatic Model Assessment for Insulin Resistance (HOMA-IR) in WT and *Dnase1l3*-KO mice after 12 weeks on ND or 45% HFD calculated as: fasting glucose (mg/dL) × fasting insulin (µU/mL) / 405. (**F**) Plasma levels of LDL cholesterol, total cholesterol, and triglycerides (TG) in WT and *Dnase1l3*-KO mice after 12 weeks on ND or 45% HFD. (**G**) Liver mass in WT and *Dnase1l3*-KO mice after 12 weeks on ND or 45% HFD. (**H–I**) Circulating alanine aminotransferase (ALAT) levels measured at 6 weeks (**H**) and 12 weeks (**I**) of ND or 45% HFD feeding. (**J**) Semi-automated quantification of hepatic steatosis in WT and *Dnase1l3*-KO mice after 12 weeks on the indicated diets. (**K**) Representative hematoxylin and eosin (H&E)-stained liver sections from WT and *Dnase1l3*-KO mice after 12 weeks on ND or 45% HFD. Scale bar = 400 μm. Data were pooled from two independent experiments with the number of mice indicated. Results are presented as mean ± SEM. Statistical analyses were performed using unpaired two-tailed t-tests and two-way ANOVA. Significance is denoted as: ns = not significant; *p ≤ 0.05; **p ≤ 0.01.

### The deficiency in DNASE1L3 exacerbates metabolic tissue inflammation induced by obesity

Given that DNASE1L3 regulates the quantity and quality of extracellular self-DNA released by dying cells (*30*), and that such self-DNA has been reported to trigger inflammatory responses during obesity (*20*), we investigated whether *Dnase1l3*-deficiency could influence adipose tissue inflammation. To address this, we analyzed the immune cell compartment (CD45⁺ cells) within the stromal vascular fraction (SVF) of the EAT from WT and *Dnase1l3*-KO mice fed either ND or 45% HFD. Flow cytometry analysis revealed a marked increase in the accumulation of CD45⁺ immune cells in the EAT of HFD-fed *Dnase1l3*-KO mice compared to HFD-fed WT controls (**Fig. 5A**). Further characterization of immune cell subsets within the EAT (**Fig. S5A**) showed that *Dnase1l3*-deficiency led to significantly higher absolute numbers of T cells, B cells, cDCs, and pDCs in HFD-fed mice relative to WT counterparts (**Fig. S5B**). Macrophages, defined by CD11b and CD64 expression, accumulated markedly in the EAT of HFD-fed *Dnase1l3*-KO mice (**Fig. 5B**). We next assessed macrophage polarization by examining the proportion of pro-inflammatory M1-*like* (CD11c⁺) and anti-inflammatory M2-*like* (CD301b⁺) macrophages (**Fig. 5C**). The frequency of CD11c⁺ M1-*like* macrophages was significantly increased in *Dnase1l3*-deficient mice following HFD exposure (**Fig. 5D**), whereas CD301b⁺ M2-*like* macrophages were reduced only in HFD-fed *Dnase1l3*-KO mice (**Fig. 5E**). Accordingly, absolute numbers of M1-*like* macrophages increased, while absolute numbers of M2-*like* macrophages remained unchanged in *Dnase1l3*-KO mice fed a HFD (**Fig. 5F-G)**. This shift toward a pro-inflammatory macrophage profile was further reflected by an increased M1/M2 ratio in *Dnase1l3*-deficient mice compared to WT mice on HFD or WT and *Dnase1l3*-KO mice on ND (**Fig. 5H**). Similar data were observed in male (**Fig. S5C-F**) and female (**Fig. S5G-J**) mice fed a 60% HFD. However, in mice fed a 60% HFD, pro-inflammatory macrophage accumulation was further exacerbated compared to those fed a 45% HFD, rendering the differences between *Dnase1l3*-KO mice and WT controls less apparent (**Fig. S5C–J**). This suggests that the pronounced immune remodeling induced by the 60% HFD masks DNASE1L3-dependent effects at the experimental endpoint. We next assessed systemic inflammation by measuring circulating cytokines and chemokines using a multiplex ELISA. The 45% HFD did not induce any detectable systemic inflammation (data not shown). In contrast, the 60% HFD led to a selective increase in circulating IL-6, IL-10, and MCP-1/CCL2 levels in *Dnase1l3* KO mice compared with WT controls (**Fig. 5I–K**). Levels of TNF-α (**Fig. 5L**) were unchanged by either diet or genotype, whereas the levels of IL-1β, IFN-α, IFN-β, and IFN-γ were undetectable in all the tested conditions (data not shown). Together, these findings indicate that *Dnase1l3* deficiency amplifies the inflammatory response that is triggered by the HFD, thereby contributing to obesity-associated metabolic complications.

**Figure 5.**
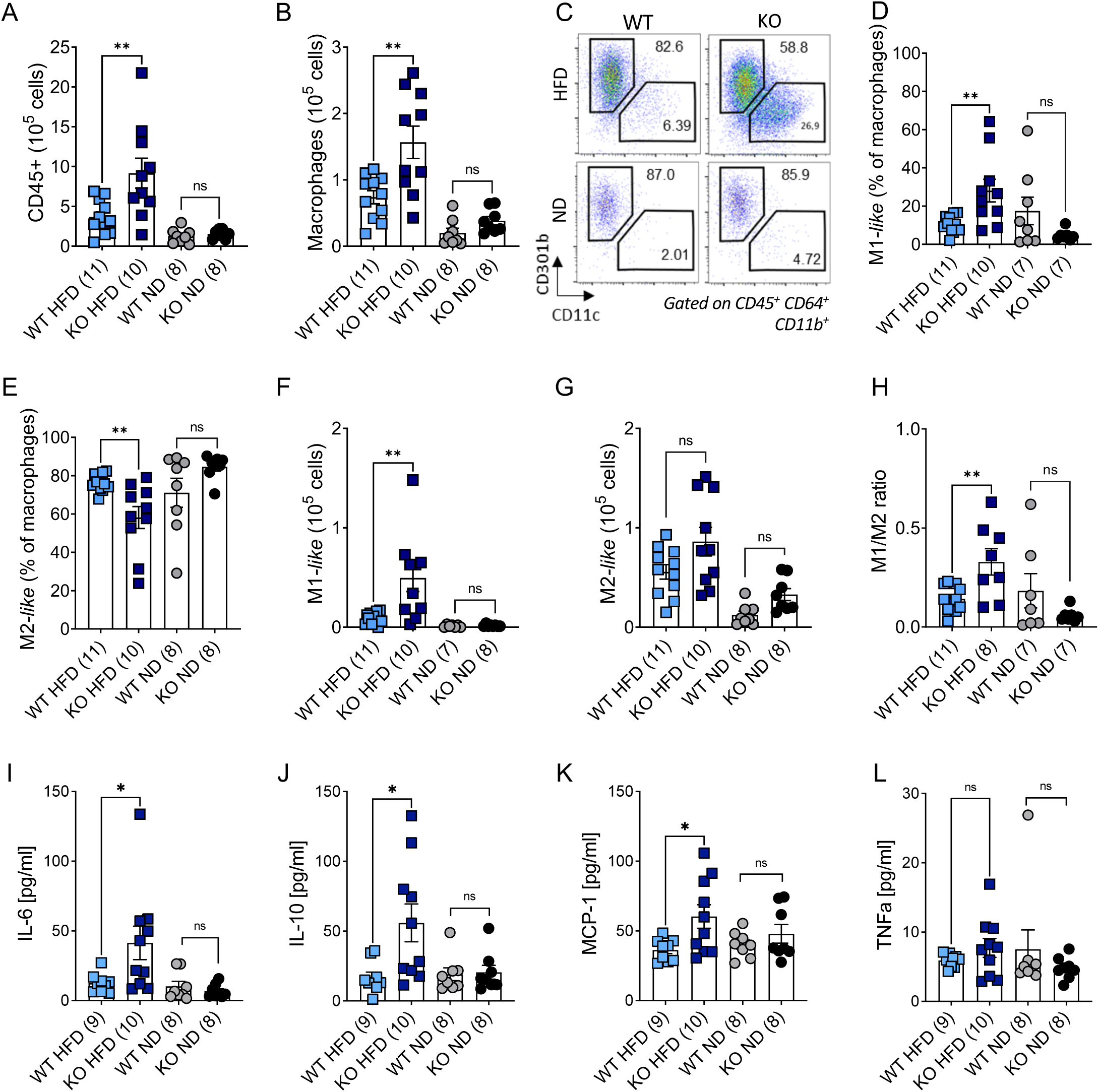
*Dnase1l3*-deficiency exacerbates metabolic tissue inflammation in 45% HFD mice. **(A)** Absolute counts of CD45^+^ cells in the stromal vascular fraction of the epididymal adipose tissue (EAT) of WT and *Dnase1l3*-KO mice on ND or 45% HFD. (**B**) Absolute counts of CD64^+^ CD11b^+^ macrophages in the EAT of WT and *Dnase1l3*-KO mice on ND or 45% HFD. (**C**) Representative flow cytometry plots of M1-*like* (CD45^+^ CD64^+^ CD11b^+^ CD11c^+^) and M2*-like* macrophages (CD45^+^ CD64^+^ CD11b^+^ CD301b^+^) in EAT. Numbers in plots indicate the frequencies of cells in each gate. (**D**) Frequency of M1-*like* macrophages (CD45^+^ CD64^+^ CD11b^+^ CD11c^+^) and (**E**) M2-*like* macrophages (CD45^+^ CD64^+^ CD11b^+^ CD301b^+^) in the EAT of WT and *Dnase1l3*-KO mice on ND or 45% HFD. (**F**) Absolute counts of M1-*like* macrophages (CD45^+^ CD64^+^ CD11b^+^ CD11c^+^) and (**G**) M2-*like* macrophages (CD45^+^ CD64^+^ CD11b^+^ CD301b^+^) in the EAT of WT and *Dnase1l3-*KO mice on ND or 45% HFD. (**H**) Ratio of the number of M1-*like* to M2-*like* macrophages in the EAT of WT and *Dnase1l3-*KO mice on ND or 45% HFD. (**I-L**). Plasma levels of IL-6, IL-10, MCP1, and TNF-α in WT and *Dnase1l3*-KO mice on ND or 60% HFD, as measured by multiplex ELISA. Data were pooled from two independent experiments with the number of mice indicated. Results are presented as mean ± SEM. Statistical analyses were performed using unpaired two-tailed t-tests. Significance is denoted as: ns = not significant; *p ≤ 0.05, **p ≤ 0.01.

### Supplementation of DNASE1L3 improves liver pathology without affecting obesity-associated weight gain or metabolic syndrome induced by HFD

Circulating DNASE activity is reduced in OB patients, and *Dnase1l3* deficiency exacerbates obesity-associated inflammation and its metabolic complications. We therefore investigated whether supplementing DNASE1L3 could hold therapeutic potential in the context of obesity. To this end, we generated AAVs encoding human DNASE1L3 (AAV-hD1L3) or luciferase (AAV-Luc). Dose titration using AAV-Luc allowed us to determine the optimal vector dose, tissue distribution, and *in vivo* persistence. Bioluminescence imaging revealed that luciferase activity persisted for at least 90 days post-injection, with the highest signal in the liver, lower levels in EAT, and minimal activity in the spleen (**Fig. S6A-C**). Consistently, human DNASE1L3 expression was predominantly detected in the liver 60 days after AAV-hD1L3 injection (**Fig. S6D**), confirming long-term and liver-specific expression. To evaluate DNASE1L3 activity, we first assessed its enzymatic function in liver lysates from *Dnase1l3*-KO mice injected with AAV-hD1L3 or AAV-Luc. AAV-hD1L3 restored nuclear DNA digestion capacity in *Dnase1l3*-KO liver lysates (**Fig. S6E**). However, this assay could not detect changes in DNASE1L3 activity in WT mice (**Fig. S6E**). Therefore, we assessed circulating DNASE activity in the serum of WT mice treated with AAV-Luc or AAV-hD1L3 and fed either ND or HFD. As observed previously, HFD reduced systemic DNASE activity, which was restored to ND levels by AAV-hD1L3 treatment (**Fig. S6F**). Notably, AAV-hD1L3 had no effect on serum DNASE activity in ND-fed mice, suggesting that DNASE1L3 activity does not exceed physiological maxima upon supplementation. Having validated the functional restoration of DNASE1L3, we next tested its therapeutic effects. WT mice were treated with AAV-hD1L3 or AAV-Luc and subsequently fed ND or 45% HFD for 12 weeks. Both groups gained similar weight upon HFD exposure (**Fig. 6A-B**) and exhibited comparable food intake (**Fig. S6G**), with no significant differences in EAT mass (**Fig. S6H**). Thus, DNASE1L3 reconstitution did not impact weight gain or adiposity. We then assessed metabolic parameters that are commonly affected by HFD. AAV-hD1L3 improved glucose tolerance and insulin sensitivity at specific time points following glucose or insulin injections (**Fig. 6C,E**). However, the area under the curve (AUC) for glucose and insulin tolerance was not significantly different between AAV-Luc- and AAV-hD1L3-treated mice (**Fig. 6D,F**). Moreover, EAT inflammation, measured by macrophage accumulation and M1-*like* polarization, remained unchanged following AAV-hD1L3 treatment (**Fig. S6I-L**). In contrast, DNASE1L3 supplementation exerted a pronounced protective effect on the liver. Although liver mass was not significantly altered by HFD (**Fig. 6G**), circulating ALAT levels were markedly increased in AAV-Luc-treated HFD mice, whereas ALAT levels in AAV-hD1L3-treated mice remained comparable to those observed under ND conditions (**Fig. 6H**). Histological analysis confirmed that AAV-hD1L3 preserved hepatic architecture and prevented HFD-induced liver injury (**Fig. 6I**). Quantitative pathological scoring revealed significant improvements following DNASE1L3 supplementation, including marked reductions in hepatic ballooning (**Fig. 6J**), steatosis (**Fig. 6K**), and inflammation (**Fig. 6L**), as evidenced by decreased immune cell infiltration (foci) across liver sections. Accordingly, the global liver pathology score of HFD-fed mice was significantly ameliorated by AAV-hD1L3 treatment (**Fig. 6M**). Notably, AAV-hD1L3 also reduced basal hepatic inflammation in ND-fed mice (**Fig. 6L**), contributing to an overall improvement in hepatic pathology under non-obesogenic conditions (**Fig. 6M**). Together, these findings indicate that AAV-mediated DNASE1L3 supplementation does not significantly influence HFD-induced weight gain or systemic metabolic dysfunction, but robustly protect against the development of MASLD. This liver-specific benefit likely reflects the strong hepatic tropism of AAV vectors and the capacity of DNASE1L3 to dampen local inflammatory responses.

**Figure 6.**
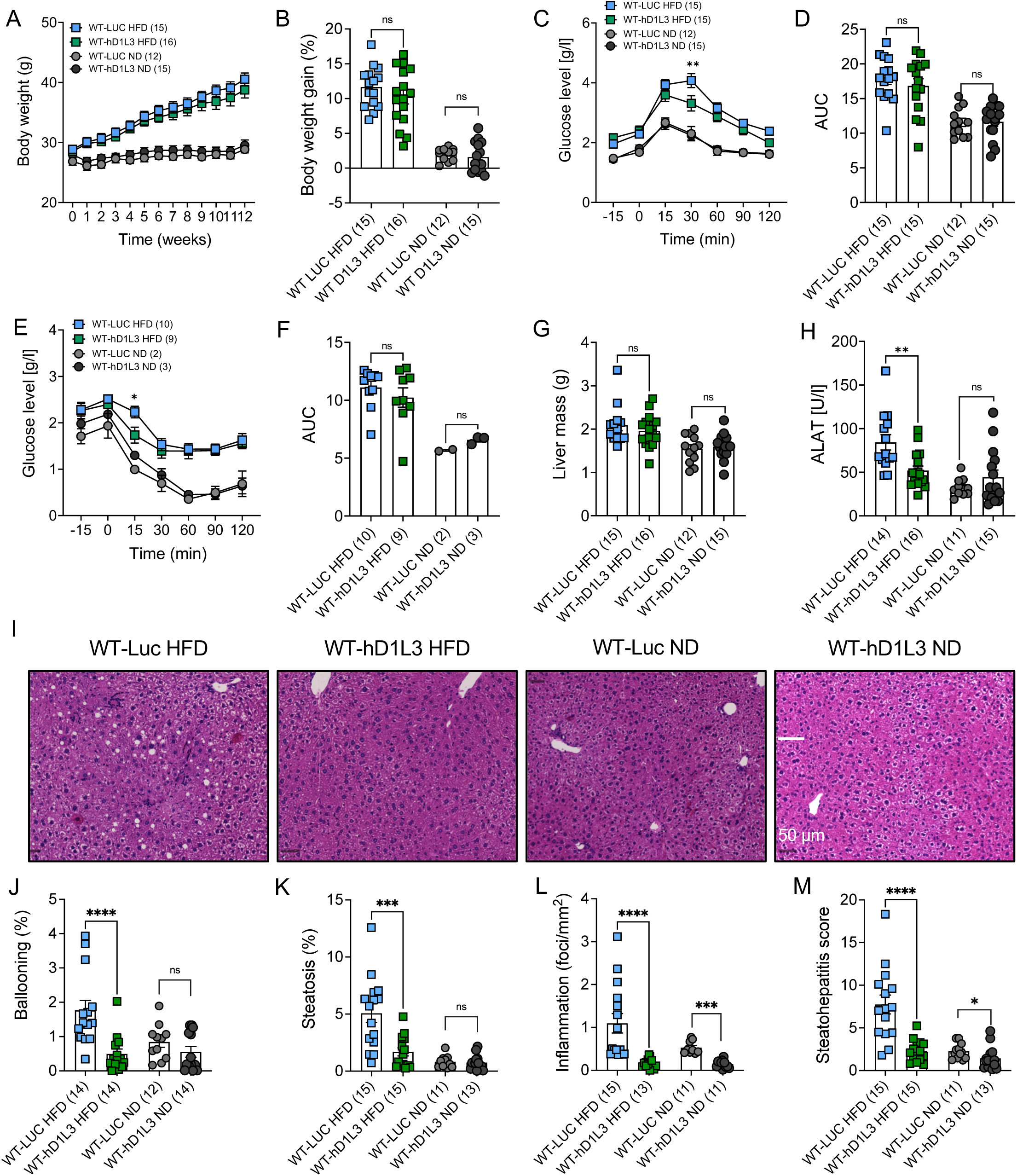
Adenoviral delivery of DNASE1L3 prevent MASLD development in HFD fed mice. WT male mice were injected with an AAV expressing luciferase (Luc) or the human DNASE1L3 (hD1L3) at a dose of 10^11^ gv/ml and exposed to ND or HFD 45% for 12 weeks. (**A**) Weekly body weight measurements and (**B**) body weight gain of the indicated mice. (**C-D**) Glucose tolerance test (GTT) (**C**) and corresponding area under the curve (AUC) (**D**). (**E-F**) Insulin tolerance test (ITT) (**E**) and corresponding area under the curve (AUC) (**F**). (**G**) Liver mass after 12 weeks of the indicated diets. (**H**) Circulating alanine aminotransferase (ALAT) levels measured at 12 weeks after diet initiation. (**I**) Representative hematoxylin and eosin (H&E)-stained liver sections after 12 weeks of the indicated diets. Scale bar = 50 μm. (**J-M**) Semi-automated quantification of liver pathology at 12 weeks of the indicated mice on the indicated diets, including hepatocyte ballooning (**J**), steatosis (**K**), inflammation (as measured by the presence of leukocyte foci/mm^2^ (**L**)) and the (**M**) overall steatohepatitis score. Data were pooled from three independent experiments with the number of mice indicated. Results are presented as mean ± SEM. Statistical analyses were performed using unpaired two-tailed t-tests or two-wat ANOVA. Significance is denoted as: ns = not significant; *p ≤ 0.05; **p ≤ 0.01; ***p ≤ 0.001; ****p ≤ 0.0001.

### DNASE1L3 supplementation restores hepatic inflammatory homeostasis in obesity

Because AAV-mediated human DNASE1L3 supplementation during obesity improved liver pathology, most notably hepatic inflammation, we next examined its impact on hepatic immune cell populations by flow cytometry. Consistent with previous reports (*8*, *9*), HFD feeding led to a marked reduction in KC (F4/80^+^, Cd11b^+^, MHC II⁺, TIM-4⁺) (**Fig. 7A,B**). Notably, AAV-hD1L3 treatment restored KC frequencies to levels comparable to those observed in ND-fed mice (**Fig. 7A,B**), consistent with a preservation of resident macrophage populations. In parallel, AAV-hD1L3 reduced total hepatic T cell infiltration (**Fig. 7C**), predominantly affecting CD8⁺ T cells (**Fig. 7D**), whereas CD4⁺ T cells were modestly increased (**Fig. 7E**). DNASE1L3 supplementation was also associated with changes in the differentiation state of liver CD8 ⁺ T cells, characterized by an increased frequency of naïve (CD44⁻CD62L⁺) and effector memory (CD44⁺CD62L⁺) cells and a concomitant reduction in effector (CD44⁺CD62L⁻) cells (**Fig. 7F–H**). Regulatory T cells (CD4⁺, FoxP3⁺) were also reduced following AAV-hD1L3 treatment (**Fig. S7A,B**). These effects on T cell abundance and phenotype were observed both in ND- and HFD-fed mice, suggesting that DNASE1L3 can influence liver immune homeostasis even in the absence of metabolic stress (**Fig. 7C–H** and **Fig. S7A,B**). These findings are consistent with histological evidence demonstrating that AAV-hD1L3 treatment attenuates hepatic inflammation in both ND- and HFD-fed mice. To further elucidate the mechanisms by which DNASE1L3 supplementation restores hepatic function, we performed 3′ bulk RNA barcoding and sequencing (BRB-seq) on livers from WT mice treated with either control AAV-Luc or AAV-hD1L3 under ND or HFD conditions. Dimensionality reduction using PCA analysis revealed that diet was the dominant driver of transcriptional variation, exerting a substantially stronger effect than AAV treatment (**Fig. 7I**). Nevertheless, PCA indicated as well that AAV-hD1L3 induced discernible transcriptional changes specifically in HFD-fed mice, whereas gene expression profiles of ND-fed mice were largely unaffected (**Fig. 7I**), suggesting that DNASE1L3-dependent effects become transcriptionally apparent primarily under metabolic stress. Consistent with this observation, AAV-hD1L3 treatment resulted in more differentially expressed genes (DEGs) under HFD than under ND conditions compared with AAV-Luc controls (916 *vs* 606 DEGs for HFD *vs* ND, respectively; **Fig. S7C,D**). As expected, HFD affected genes involved cholesterol and lipid metabolism, confirming the metabolic stress imposed by HFD (**Fig. S7C-F**). DNASE1L3 supplementation affected a similar number of genes under ND (392 DEGs) and HFD (328 DEGs) conditions; however, the magnitude of gene downregulation was greater under HFD (**Fig. 7J,K**). Across both ND and HFD conditions, AAV-hD1L3 primarily repressed immune-related gene expression (**Fig. 7J-M**). By contrast, DNASE1L3 induced genes did not segregate into coherent transcriptional programs in either dietary context (**Fig. 7J,K and data not shown**). Gene ontology analyses further confirmed that HFD robustly downregulated pathways associated with cholesterol and lipid biosynthesis while upregulating pathways related to lipid storage, oxidation and catabolism (**Fig. S7E,F**). DNASE1L3 supplementation selectively reduced pathways associated with immune and inflammatory functions in mice fed a ND or HFD (**Fig. 7L**). Notably, under ND the supplementation of human DNASE1L3 specifically reduced acute inflammatory response (*Saa1, Saa2, Saa3 and Orm2*) (**Fig. 7J,L**) and under HFD, DNASE1L3 specifically suppressed pathways linked to interferon responses and antigen presentation (**Fig. 7K,L**). Analysis of individual genes within these pathways confirmed that DNASE1L3 supplementation in obese mice limited the expression of interferon-stimulated genes (e.g. *Cxcl9, Ccl5, Ifi47, Igtp, Ly6a, Gbp6, and Ubd*), antigen presentation machinery (*H2-Aa, H2-Ab1, H2-Eb1, H2-Dma, and Cd74*), adaptive immune response markers (*Jchain, Serpinb, Cd3d, Nkg7 and Cd40*), and, to a lesser extent, macrophage activation markers (e.g. *Cd68, Mpeg1, Laptm5, Sirpa, Lst1, Axl*) (**Fig. 7M**). This immune dampening was accompanied by reduced expression of lipid biosynthesis-associated genes, as well as extracellular matrix-associated genes (*Col1a1, Col1a2, Col3a1, and Col12a1*) (**Fig. S7E,F**), which are commonly upregulated during MASLD progression and contribute to fibrotic remodeling. Collectively, these findings indicate that DNASE1L3 contributes to reduced basal immune activation under homeostatic conditions and, under metabolic stress, attenuates interferon signaling, antigen presentation, and fibrogenic gene programs in the liver, thereby limiting MASLD pathogenesis.

**Figure 7.**
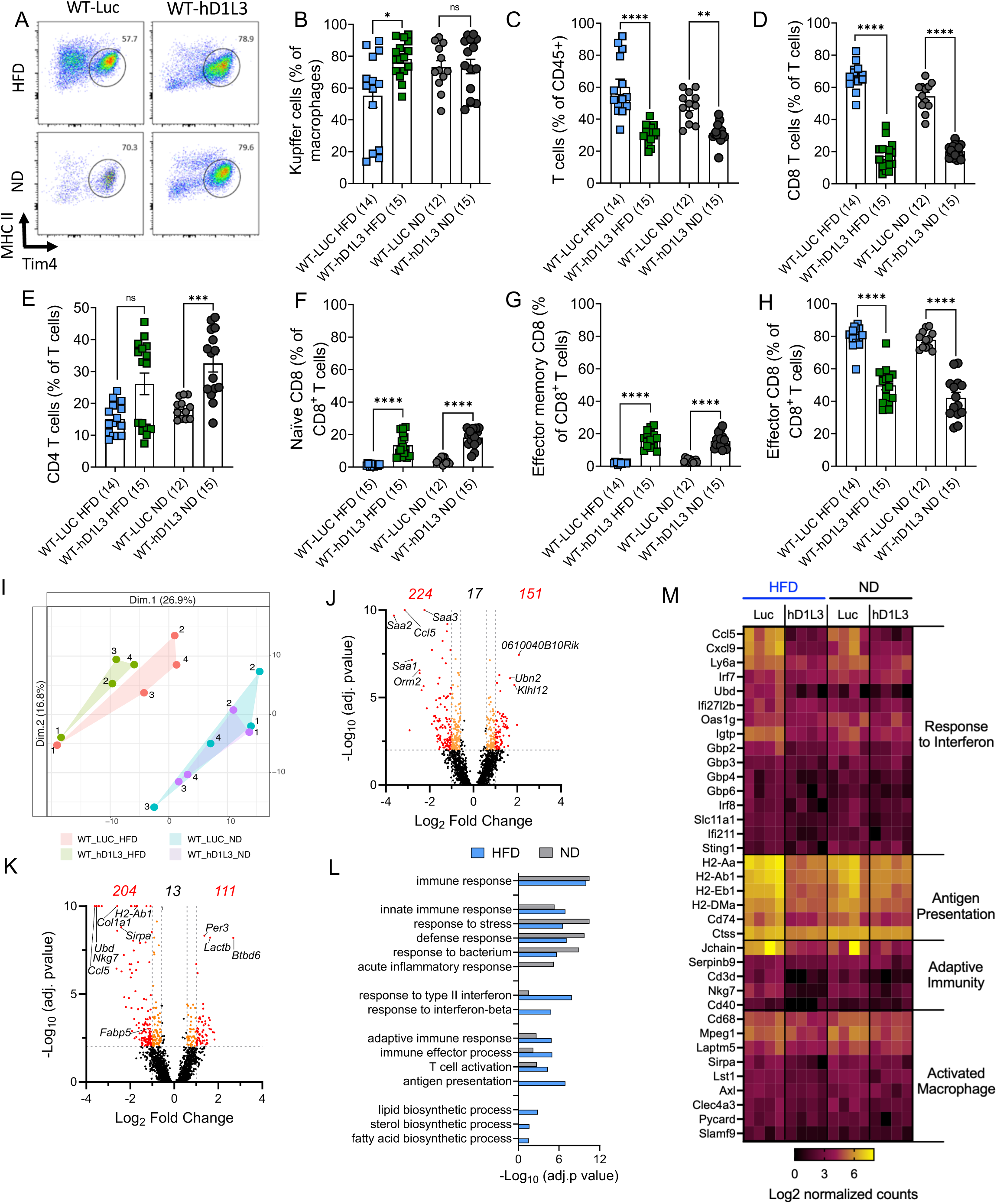
Adenoviral delivery of DNASE1L3 limits liver inflammation induced by HFD. WT male mice were injected with an AAV expressing luciferase (Luc) or the human DNASE1L3 (hD1L3) at a dose of 10^11^ gv/ml and exposed ND or HFD 45% for 12 weeks. (**A**) Representative flow cytometry plots of Kupffer cells (CD45^+^ CD11b^+^ F4/80^+^ Tim4^+^ MHCII^+^) in the liver after 12 weeks of the indicated diets and treatments. Numbers in plots indicate the frequencies of cells in each gate among CD45^+^, TCRβ^-^, CD19^-^, CD11b^+^, F4/80^+^ cells. (**B**) Frequency of Kupffer cells (CD45^+^ CD11b^+^ F4/80^+^ Tim4^+^ MHCII^+^) in the liver after 12 weeks of the indicated diets and treatments. (**C**) Frequency of T cells (CD45^+^, TCRβ^+^) in the liver after 12 weeks of the indicated diets and treatments. (**D**) Frequency of CD8 (CD45^+^, TCRβ^+^ CD8^+^) and (**E**) CD4 (CD45^+^, TCRβ^+^ CD4^+^) T cells in the liver after 12 weeks of the indicated diets and treatments. (**F**) Frequency of naïve (CD62L^+^, CD44^-^), (**G**) effector memory (CD62L^+^, CD44^+^) and (**H**) effector (CD62L^-^, CD44^+^) CD8 T cells in the liver after 12 weeks of the indicated diets and treatments. Data were pooled from three independent experiments with the number of mice indicated. Results are presented as mean ± SEM. Statistical analyses were performed using unpaired two-tailed t-tests or two-wat ANOVA. Significance is denoted as: ns = not significant; *p ≤ 0.05; **p ≤ 0.01; ***p ≤ 0.001; ****p ≤ 0.0001. (**I**) PCA of BRB-seq data from livers of WT mice treated with AAV-Luc or AAV-hD1L3 and fed either ND or HFD for 12 weeks. Volcano plots identifying differentially expressed genes from the liver from WT mice treated with AAV-hD1L3 relative to WT mice treated with AAV-Luc and fed either a ND (**J**) or a HFD (**K**). (**L**) Gene ontology (GO) analysis of the downregulated genes of the liver from WT mice treated with AAV-hD1L3 compared to AAV-Luc on ND and HFD. (**M**) Heat map showing log2 normalized expression of the indicated genes in individual mice treated with AAV-Luc or AAV-hD1L3 and fed either ND or HFD for 12 weeks.

## DISCUSSION

Emerging evidence implicates extracellular self-DNA and its sensing by innate immune receptors in the pathogenesis of obesity (*20*, *21*). Here, we identify DNASE1L3 as a critical regulator of cfDNA homeostasis that protects against obesity-associated metaflammation and metabolic dysfunction.

We observed elevated levels of cfDNA in OB individuals, consistent with previous reports (*22*, *51*). Using a qPCR-based approach, we determined that this cfDNA was primarily of nuclear origin and soluble. Although the total number of circulating MPs was reduced in OB patients, the DNA content per MP was significantly higher, suggesting an accumulation of MP-associated cfDNA, which is an extracellular DNA pool regulated by DNASE1L3 (*33*). While earlier studies reported increased levels of MPs in OB individuals (*52*, *53*), those were primarily derived from platelets, which lack nuclei and were excluded from our analysis. The observed increase in cfDNA correlated positively with key metabolic parameters, including BMI, insulin levels, and HOMA-IR, in line with previous findings (*22*, *51*). Furthermore, we found that bariatric surgery led to a significant reduction in cfDNA levels. While reinforcing the link between cfDNA burden and metabolic status, this result contrasts with earlier studies (*51*), likely due to the higher sensitivity of our cfDNA detection method and the longer post-operative follow-up (>6 months) of our cohort. Although we could not pinpoint the cellular origin of cfDNA in OB individuals, its levels correlated with neutrophil-derived proteins (MPO, NE), implicating NETs. This aligns with recent findings of increased circulating MPO-DNA complexes in severe obesity (*41*, *54*). While our study focused on systemic changes, cfDNA may also accumulate in metabolic tissues (*25*, *55*), further amplifying inflammation.

We also found qualitative changes in cfDNA in obesity, including increased fragment size and altered end-motifs, indicative of impaired DNASE activity (*56*). Two main DNASEs, including DNASE1 and DNASE1L3, contribute to total circulating DNASE activity (*49*). DNASE1L3 (*43*, *44*) but not DNASE1 (*57*), shapes cfDNA fragmentation and end-motifs suggesting that cfDNA abnormalities observed in obesity are likely due to impaired DNASE1L3 activity. Accordingly, OB individuals showed abnormalities in cfDNA properties resembling those of D1L3^null^ patients. Although DNASE1L3 expression was not reduced, total circulating DNASE activity was impaired in both OB individuals and HFD-fed mice, suggesting extracellular or post-translational inhibition. Given prior reports that DNASE1L3 function can be blocked by circulating autoantibodies in systemic autoimmune disease (*47*, *48*), we investigated this possibility in obesity. We identified anti-DNASE1L3 autoantibodies in a subset of OB patients in two independent cohorts, while anti-DNASE1 antibodies were absent. Although the limited prevalence of these antibodies precludes strong correlative conclusions, these findings suggest that obesity may promote B cell tolerance breakdown, as previously described (*24*, *39*), and impair DNASE1L3 function through autoimmune mechanisms. Additionally, metabolic factors such as dyslipidemia may further inhibit circulating DNASE activity (*38*).

To directly assess the role of DNASE1L3 in obesity, we used *Dnase1l3*-deficient mice fed with HFD. These mice developed greater weight gain, glucose intolerance, insulin resistance, metaflammation, and MASLD compared WT controls. *Dnase1l3* deficiency altered feeding rhythmicity without affecting overall caloric intake and was accompanied by modest changes in energy expenditure, potentially increasing susceptibility to HFD. In addition, DNASE1L3 is highly expressed in innate immune cells along the intestinal epithelium (*58*, *59*); thus, its deficiency could enhance nutrient absorption and promote weight gain independently of overall caloric intake or energy expenditure. The increased weight gain in *Dnase1l3*-KO mice was consistent across both 45% and 60% HFDs and in both sexes, supporting a robust, sex-independent role for DNASE1L3 in obesity. Under a 60% HFD, the metabolic impact of *Dnase1l3* deficiency was more pronounced at 6 weeks than at 12 weeks, likely reflecting the severe and rapidly progressive metabolic stress imposed by such diet (*60*, *61*). In contrast, on 45% HFD, which is more representative of typical Western diets (*61*), metabolic disease developed progressively, becoming evident only after 12 weeks *in Dnase1l3*-KO mice. Thus, *Dnase1l3*-deficiency appears to accelerate the onset or heighten susceptibility to obesity-associated complications. Even though female mice were relatively protected from metabolic disease induced by HFD as previously reported (*50*), *Dnase1l3*-deficiency still increased EAT inflammation and induced MASLD, highlighting its sex-independent role.

*Dnase1l3*-deficient mice on HFD showed enhanced systemic inflammation and immune cell infiltration in the EAT, particularly of pro-inflammatory M1-*like* macrophages, which are key mediators of obesity-driven metaflammation and the ensuing health complications (*5–7*). We propose DNASE1L3 restrains inflammatory responses by limiting the abundance and immunostimulatory potential of extracellular self-DNA. Although we did not directly test pathways involved in DNA sensing, such as TLR9 or cGAS/STING, in the exacerbation of obesity-induced metabolic inflammation induced by the loss of DNASE1L3, prior research supports this model (*22–29*). Our data position DNASE1L3 upstream of these pathways, regulating cfDNA burden and immunostimulatory potential.

Recombinant DNASE1 treatment in HFD-fed mice has previously shown limited therapeutic efficacy (*24*), possibly due to its inability to digest complex DNA structures (*31*). Given that DNASE1L3 has broader nuclease activity (*31*, *33*), we tested whether its overexpression could mitigate obesity-driven inflammation and pathology. AAV-mediated delivery of human DNASE1L3 prevented MASLD development, reduced liver inflammation and T cell infiltration, without significantly affecting body weight or EAT inflammation. This tissue-specific protection likely reflects the hepatic tropism of the AAV and the importance of DNASE1L3 function in liver-resident immune and endothelial cells (*45*). Notably, *DNASE1L3* overexpression restrained liver inflammation even in the absence of HFD, suggesting a homeostatic role for DNASE1L3 in maintaining liver immune balance. Under HFD, DNASE1L3 more selectively inhibited the expression of IFN- and antigen presentation–related transcriptional program. By limiting the immunostimulatory potential of extracellular self-DNA, particularly its ability to trigger IFN responses, we believe that DNASE1L3 dampens T cell activation, an essential driver of MASLD pathogenesis (*62*, *63*). Emerging data also implicate DNASE1L3 in limiting MASLD evolution to hepatocellular carcinoma (*64*, *65*), and our findings extend its protective role to MASLD development.

In conclusion, obesity impairs DNASE1L3 activity, leading to cfDNA accumulation altered fragmentation that contribute to metaflammation and metabolic disease. DNASE1L3 emerges as a critical regulator of cfDNA homeostasis and an attractive target for mitigating obesity-related complications, particularly in the liver.

## MATERIALS and METHODS

### Sex a biological variable

Our human cohort included both male and female participants, and both sexes were also represented in the mouse experiments. Given the known influence of sex on body weight and metabolic outcomes, HFD-induced weight gain and metabolic complications were analyzed separately by sex.

### Key Resources

All the key resources, their origin, their specification as well as their concentration of usage are indicated in the **Supplementary Table 1.**

### Mice and diets

*Dnase1l3*-deficient (*Dnase1l3*^LacZ/LacZ^) mice, generated by targeted germline replacement (model TF2732, Taconic Knockout Repository), were backcrossed for over 10 generations onto the C57BL/6J background. Homozygous KO animals were obtained through intercrossing. Age-matched WT C57BL/6J mice, bred in the same colony, were used as controls. All mice were maintained in a specific-pathogen-free facility at the University of Bordeaux (Animalerie A2, approval #B33-063-91) and cohoused prior to the start of experiments to minimize microbiota-driven variation. Mice (12–15 weeks old) were housed under controlled temperature and humidity with a 12-hour light/dark cycle and had ad libitum access to water and food. Animals were fed either a standard rodent chow (A03 or AIN93G AMF butter, SAFE, Augy, France) or a HFD containing 45% or 60% fat (246HF or 260HF, SAFE) for 12 weeks. When indicated, mice were also placed in individual metabolic cages (24 phenomaster cages, TSE Systems GmbH, Bad Homburg, Germany) where locomotor activity, food intake and gas exchanges were measured under controlled conditions of illumination, temperature and humidity as previously described (*66*). The number and sex of mice used in each experiment are detailed in the main text and/or figure legends. All animal procedures complied with the European Directive 2010/63/EU for the protection of animals and were approved by the French Ministry of Research (protocols #20125 and #33484).

### Human Subjects and tissue samples

Included OB patients (n=67) were aged between 22 and 65 years and followed in the Nutrition Department of the Bordeaux University Hospital or the services of digestive and bariatric surgery departments of two private clinics from Bordeaux area (Tivoli-Ducos and Jean-Villar). Patients that underwent bariatric surgery had severe obesity meeting the eligibility criteria for surgery (according to the guidelines of the French Health Authority: BMI ≥40 kg/m2 or ≥35 kg/m2 in the presence of complications susceptible to be improved by bariatric surgery). The exclusion criteria of patients that underwent bariatric surgery were: age > 65 years old; acute or chronic inflammatory conditions (other than obesity or obesity-related comorbidities); current treatment with antidepressants or any other psychotropic drug; current treatment with anti-inflammatory or immunosuppressive drugs; current diagnosis of psychiatric disease (except for major depressive disorder); and/or severe medical illness. The study was approved by the local Committee for the Protection of Persons (Bordeaux, France). All patients provided written informed consent after reading a complete description of the study. Anthropometric and biological data were collected under conditions of regular clinical care. Blood from sex- and age-matched healthy donors (n=43) (BMI <30 kg/m2) were obtained from the local Blood Transfusion Center (Etablissement Français du Sang - EFS) (**Table 1-2**). Greater omental adipose tissue samples were collected from OB patients undergoing bariatric surgery (n=22) and lean patients undergoing gastric surgery for non-obesity related indications (n=10) were recruited at ILS hospital Salt Lake, Kolkata, India. All participants provided written informed consent in accordance with the recommendations of the institutional review boards of the participating institutions. To further validate the presence of anti-DNASE1L3 antibodies in OB individuals, we included an additional cohort provided by the Biobank of the Aragon Health System (National Registry of Biobanks B. B.0000873, PT23/00146), integrated into the ISCIII Biobanks and Biomodels Platform (**Table 3**). All samples were processed according to standardized operating procedures and with approval from the corresponding Ethics and Scientific Committees. Finally, two patients with pediatric hypocomplementemic urticarial vasculitis syndrome carrying a homozygous *DNASE1L3* variant (c290-291del) were also included in the study. Patients were diagnosed and treated at the Pediatric Clinical Immunology Department, Hôpital Pellegrin, Bordeaux, France and the sample collection of patients was approved by the Institutional Ethical Committee.

### Adeno-Associated Viruses

Adeno-associated viral vectors encoding human DNASE1L3 or Luciferase were produced in HEK293 cell cultures and purified by iodixanol density gradient ultracentrifugation by the Vect’Ub facility at the University of Bordeaux. The serotype pAAV2/9n used in our study was a gift from James M. Wilson (Addgene plasmid # 112865). AAVs were injected i.v at 1×10^11^ gv/ml per mouse. After 2 weeks and up to 12 weeks, Luciferase activity and human *DNASE1L3* expression were validated by bioluminescence imaging and RT-qPCR respectively.

### Murine body weight, food intake and feeding efficiency measurements

Individual mouse body weight and food intake per cage were recorded weekly during all the experiments. 12 weeks after HFD initiation, mice were isolated in single cage to measure individual food intake during 8 days. Feeding efficiency was calculated as the ratio between cumulative body weight gain and cumulative caloric intake over the same period of time.

### Body composition analysis

Mouse fat and lean body composition analysis was performed with the Echo Magnetic Resonance Imagery (MRI) (EchoMedical Systems, Houston, TX, USA) 12 weeks after the initiation of the dietary intervention.

### Glucose and insulin tolerance test

At baseline, 6 and 12 weeks following initiation of the HFD, mice were subjected to either a GTT or ITT. For GTT, mice received an i.p. injection of 10% D-(+)-glucose at a dose of 1 g/kg body weight. For ITT, mice were injected i.p. with insulin at 0.75 U/kg body weight. Prior to testing, animals were fasted for 6 hours. Blood glucose levels were measured from the tail vein using an Accu-Chek Performa glucometer (Roche Diabetes Care, France) at baseline (0 min) and at 15, 30, 60, 90, and 120 minutes following glucose or insulin injection.

### Murine plasma collection and analysis

Blood samples were collected by submandibular puncture at baseline, as well as 6 and 12 weeks after HFD initiation. Samples were drawn into EDTA-coated tubes, centrifuged at 500 × g for 10 minutes at 4°C, and the resulting plasma was stored at - 80°C until analysis. Plasma levels of total cholesterol, LDL, TG, and ALAT were measured using a Pentra C400 automated clinical chemistry analyzer (Horiba, France). Plasma insulin concentrations were quantified by ELISA (Crystal Chem Inc, #90080). Insulin sensitivity was assessed using the Homeostasis Model Assessment of Insulin Resistance (HOMA-IR), calculated as: HOMA-IR = (fasting insulin [µU/ml] × fasting glucose [g/l]) / 405. Plasma levels of inflammatory cytokines were quantified using the MesoScale Discovery U-PLEX murine proinflammatory panel 1 as per the manufacturer’s instructions.

### Murine tissue collection, processing and immune cell isolation

EAT, IAT and liver were collected, weighed, and immediately placed in cold media. AT samples were digested in PBS containing 0.5% bovine serum albumin (BSA), 8 mg/mL collagenase type I-A (Sigma), and 10 mM CaCl₂ for 20 minutes at 37°C. The resulting cell suspension was filtered through a 70 μm strainer and centrifuged at 450 × g for 5 minutes at room temperature (RT). The cell pellet, corresponding to the SVF, underwent red blood cell lysis, was washed in PBS, and resuspended in FACS buffer (PBS with 1% FBS and 2 mM EDTA). Livers were perfused through the inferior vena cava with PBS, weighed, and processed into single-cell suspensions following previously described protocols (*67*). CD45⁺ immune cells were enriched by magnetic-activated cell sorting using CD45 microbeads (Miltenyi Biotec). Cell numbers were determined using a BD Accuri™ C6 flow cytometer (BD Biosciences).

### Human blood samples processing

Blood was collected in 7 ml EDTA tubes and rapidly spun at 3,500 × *g* for 15 min without brakes. Platelet-poor plasma was gently collected and spun again at 3,500 × *g* for 15 min without brakes to obtain platelet-free plasma (PFP). PFP were stored at -80°C. PBMCs were isolated from the remaining blood by Ficoll density gradient centrifugation. Absolute PBMCs counts were determined by the BD-Accuri™ C6 flow cytometer (BD-Biosciences), and used directly for flow cytometry.

### Flow cytometry staining, sorting and acquisition

Mouse single-cell suspensions were first incubated with anti-CD16/32 Fc-blocking antibody for 10 minutes at 4 °C to prevent nonspecific antibody binding. Following blocking, cells were stained with a fixable viability dye (Zombie Aqua or ViaDye™ Red; BioLegend) to exclude dead cells. After washing in FACS buffer (PBS supplemented with 1% FBS and 2 mM EDTA), cells were incubated with extracellular antibodies (listed below and in **Supplementary Table 1**). The extracellular antibody panel included: anti-CD45 PerCP-Cy5.5, anti-TCR-β APC-eFluor 780, anti-B220 PE-CF584, anti-B220 BV421, anti-B220 PE-Dazzle, anti-CD19 BUV737, anti-CD8 APC, anti-CD8 BV785, anti-CD4 FITC, anti-CD4 BUV494, anti-CD44 BV510, anti-CD62L BUV395, anti-Ly6C Pacific Blue, anti-Ly6C BV570, anti-F4/80 FITC, anti-F4/80 BV650, anti-CD11b APC-Cy7, anti-CD11b redFluor 710, anti-CD64 BV605, anti-CD301b PE-Cy7, anti-TIM4 APC, anti-ESAM PE, anti-CD206 BV711, anti-MHCII Pacific Blue, anti-CD11c APC-Cy7, anti-CD11c BUV805, anti-CD11c PE, anti-CD317 BV711, anti-Siglec H BV421, anti-Ly6G BV711, and anti-Ly6G Spark-NIR 685. After surface staining, cells were fixed using the FoxP3 Fixation/Permeabilization kit (Fisher Scientific) according to the manufacturer’s instructions. Intracellular staining was then performed using anti-FoxP3 PE or anti-FoxP3 PE-Cy5 antibodies in FoxP3/Transcription Factor Staining Buffer (Fisher Scientific) for 45 minutes at RT. Samples were acquired using either a BD LSRFortessa™ (BD Biosciences) or Cytek Aurora (Cytek) flow cytometer, depending on the experiment. Data were analyzed using FlowJo software version 10 (BD Biosciences). In humans, pDCs, DCs and CD14^+^ cells from VAT were sorted as previously described (*25*).

### Histological analysis of murine metabolic tissues

Murine EAT and liver samples were fixed in 10% neutral-buffered formalin (Sigma, HT501128-4L) for 24 h at RT, then stored in 70% ethanol and embedded in paraffin. Tissue sections were stained with hematoxylin and eosin and imaged at 40 X magnification (Histology Facility, Necker Hospital, Paris). Adipocyte surface area and perimeter were quantified using NDP.view2 Plus software (Hamamatsu Photonics, France). Hepatic steatosis was assessed blindly to genotype and diets by a trained pathologist using QuPath bioimage analysis software (*68*), with segmentation performed via the SLIC (Simple Linear Iterative Clustering) tool (*69*).

### Isolation of microparticles

PFP was centrifuged at 22,000 × g for 1 hour at 4 °C to pellet MPs. The resulting supernatant was collected as the soluble, MP-depleted fraction (MP-), while the pellet was resuspended to obtain the MP-enriched fraction (MP+). To quantify MPs, 1 µl of the MP+ fraction was stained with anti-human CD235a-PE (to exclude red blood cell–derived vesicles) and CD41-APC (to exclude platelet-derived vesicles). The double-negative population was quantified using a BD Accuri™ C6 Plus flow cytometer (BD Biosciences). A forward scatter (FSC) threshold of 2500 was applied to exclude events smaller than MPs and reduce background noise

### Quantification of cfDNA

#### Human cfDNA

The cfDNA was purified from the whole plasma (WP), MP- and MP+ fractions using a QIAamp DNA Blood Mini Kit (QUIAGEN). Nuclear DNA concentration was quantified by using Alu primers that targeted Alu repetitive elements of 200bp (FW: AAAATTAGCCGGGCGTG, RV: AGACGGAGTCTCGCTCTGTC). All qPCRs were performed using GoTaq® qPCR Master mix (Promega) on a CFX384 thermocycler (Bio-Rad TM). The cycling conditions were: initial denaturation of 12 min at 95°C, followed by 45 amplification cycles of denaturation at 95°C for 20 sec, annealing at 56°C for 1 min, and elongation at 72°C for 1 min. cfDNA levels were calculated according to a standard in each fraction. Total plasma concentration of mitochondrial DNA in each sample was measured using amplification of a 67 bp MT-CO3 gene sequence as previously described (*70*). qPCR was conducted in a 25 μl reaction volume on a CFX96 instrument with CFX management software 3.0 (Bio-Rad). Thermal cycling involved a 3-minute Hot-start Polymerase activation denaturation step at 95°C, followed by 40 cycles at 95°C for 10 s and 60°C for 30 s. Sample concentrations were extrapolated using the standard curve. All experiments included duplicate samples and DNA quantification data were validated using an internal control with a known concentration. The mitochondrial DNA to nuclear DNA ratio (MNR) was expressed as the ratio of the mean of mitochondrial DNA copy nb/ml of plasma value to the mean of nuclear DNA copy nb/ml of plasma value.

#### Mouse cfDNA

Nuclear and mitochondrial DNA concentration was quantified by using intra-B1 (FW: GGGCATGGTGGCGCACGCCT, RV: GAGACAGGGTTTCTCTGTGT and mtND1 (FW: CTAGCAGAAACAAACCGGGC, RV: CCGGCTGCGTATTCTACGTT) primers, respectively. qPCR method have been reported previously (*71*). Briefly, reaction was subjected to an initial denaturation of 8 min at 95°C, followed by 40 amplification cycles of denaturation for 30s at 95°C, annealing for 40s at 55°C, and elongation for 1 min at 72°C.

### DNA library preparation for soluble whole genome sequencing (sWGS)

DSP (double-strand protocol) libraries were prepared with the NEB Next® Ultra™ II kit. A minimum of 2 ng of cirDNA was engaged without fragmentation, and kit provider’s recommendations were followed. Illumina paired-end adaptor oligonucleotides are ligated on repaired A-tailed fragments, then purified by solid-phase reversible immobilization (SPRI) and enriched by 11 PCR cycles with UDI primers indexing, then SPRI purified again. SPRI purification was adjusted to keep the small fragments around 70 bp of insert. Finally, the libraries to be sequenced were precisely quantified by qPCR, in order to load the appropriate DNA quantity to the Illumina sequencer, and to obtain a minimum of 1.5 million clusters.

### Fragment size and end motifs extraction from sWGS data

All libraries were sequenced on MiSeq 500 or NovaSeq (Illumina) as Paired-end 100 bp reads. Image analysis and base calling were performed using Illumina Real Time Analysis with default parameters. The individual barcoded paired-end reads were trimmed with Cutadapt v1.10, to remove the adapters and discard trimmed reads shorter than 20 pb. Trimmed FASTQ files were aligned to the human reference genome (GRCH38), using the MEM algorithm in Burrows-Wheeler Aligner v0.7.15. Supplementary alignments, PCR duplicates, and reads with MapQ < 30 were filtered out from the analysis. Coverage values (mean±sd) were assessed for both the DSP and SSP for all sample groups. For the DSP, the mean coverage was 0.52 ± 0.24. The insert sizes (=fragment sizes) were then extracted from the aligned bam files with the TLEN column for all pairs of reads having an insert size between 0 and 1,000 bp. The frequency of each fragment size was then normalized to the total fragment count to generate relative size distributions.We determined the end motif sequence of the cirDNA fragments based on the filtered BAM files aligned against the reference human genome GRCh38. End motifs were determined for 5’ and 3’ DNA fragment ends. The sequences of end motifs were extracted and compiled using the pysam and pandas Python packages. For fragments mapped on the positive strand, 5’ end motifs were extracted as the n first nucleotides, and 3’ end motifs as the n last nucleotides (n being the length of the end motifs analyzed, i.e., 3, 2 or 1 bp). For fragments mapped on the negative strand, 5’ end motifs were extracted as the reverse complement of the n first nucleotides, and 3’ end motifs as the reverse complement of n last nucleotides. The relative frequency of each motif within each individual sample (or the fragment size fraction of the sample) was computed. When comparing 5’ and 3’ end motifs frequencies, the reverse complement of 5’ end motifs were used. Machine learning was performed using R (v 4.3.3) and the software packages ggplot2 (3.5.1) for plotting, and built-in R function prcomp() for PCA.

### Detection of Anti-DNASE1 and anti-DNASE1L3 antibodies by ELISA

Nunc™ MaxiSorp™ plates were coated overnight at 4 °C with 50 µl/well of 2 µg/ml recombinant DNASE1 (Roche) or purified recombinant DNASE1L3 in PBS, as previously described (*46*). After coating, plates were washed with PBS containing 0.05% Tween-20 and 1% nonfat dry milk (PBS-T/1% NFDM), then blocked with 250 µl/well of PBS containing 4% NFDM for 6 hours at RT. Following three washes, plasma samples diluted 1:50 in PBS were added and incubated overnight at 4 °C. Plates were then washed and incubated for 1 hour at RT with goat anti-human IgG-HRP (Ozyme, BETA80-104P) diluted 1:150,000 in PBS with 1% NFDM. After three final washes, plates were developed with TMB substrate (Thermo Fisher Scientific) and the reaction was stopped with H₂SO₄. Absorbance was measured at 450 nm using a CLARIOstar™ plate reader (BMG Labtech). Background signals from each plasma were measured in parallel and subtracted from the final readings.

### DNASEs activity measurement

#### SRED assay

DNASE activity was assessed using the Single Radial Enzyme Diffusion (SRED) assay. The gel solution was prepared by combining distilled water with calf thymus double-stranded DNA (1 mg/mL), Tris-HCl (20 mM, pH 7.8), MnCl₂ (10 mM), CaCl₂ (2 mM), and ClearSight™ DNA stain. This mixture was heated at 50 °C for 10 minutes, then mixed 1:1 with distilled water containing 2% ultrapure agarose. The final solution was poured into a casting plate and allowed to solidify at RT. After solidification, 1.5 mm wells were punched at 2 cm intervals using a biopsy punch. Recombinant DNASE1 and 10μl of plasma sample were deposited into each well, and gels were incubated at 37 °C for 16 hours. DNASE activity was visualized under UV light as dark halos, the diameter of which reflected enzymatic activity.

#### PicoGreen™ assay

To quantify DNASE activity with higher sensitivity and throughput, a fluorescence-based assay using the intercalating dye PicoGreen™ was employed. Briefly, 50 ng of calf thymus dsDNA in Tris-HCl buffer was incubated with plasma samples at a 1:1 ratio for 2 hours at 37 °C. Enzymatic digestion was then halted by adding Tris-EDTA containing PicoGreen™ (1:200 dilution). Fluorescence, inversely proportional to DNASE activity, was measured using a CLARIOstar™ plate reader (BMG Labtech). The DNA content of plasma was independently quantified using PicoGreen™ and subtracted from the experimental readings. Results were expressed as the % of DNA degradation, using undigested DNA (50 ng) as a reference.

### RNA-seq library preparation and data analysis

Total RNA was extracted from liver using the RNeasy Mini Kit (Qiagen), following the manufacturer’s instructions. RNA concentration and purity were assessed with a NanoDrop™ ND-1000 spectrophotometer (Thermo Fisher Scientific), and only samples with RNA Integrity Number (RIN) ≥ 7 were used for library preparation. 3′ BRB-seq was performed as previously described (*72*). Reverse transcription and template switching were performed using 2.5 ng/µL of total RNA. Double-stranded cDNA was synthesized by PCR, and 50 ng was tagmented using the Illumina Nextera XT Kit. Libraries were quantified with a Quantus™ fluorometer (Promega) and sequenced on an Illumina NovaSeq X (Macrogen). Reads were demultiplexed using sample-specific 6-bp barcodes and aligned to the mm10 reference transcriptome using BWA (v0.7.17) after trimming with Cutadapt (v3.5). Gene expression was quantified via 10-bp UMI counts. Low-quality samples (<25,000 reads or <8000 genes detected) and underpowered conditions were excluded. Lowly expressed genes were filtered out. Differential expression was analyzed with DESeq2 and DEGreport, using a Wald test with Benjamini–Hochberg correction (p < 0.05; fold change ≥ 1.5).

### Statistical analysis

Statistical analyses were conducted using GraphPad Prism version 9.4.1. The normality of data distribution was first evaluated using the D’Agostino & Pearson omnibus normality test. For normally distributed data, parametric tests were applied, including unpaired t-tests or two-way ANOVA as appropriate. For non-normally distributed data, non-parametric tests such as the Mann–Whitney U test or the Kruskal–Wallis test followed by Dunn’s multiple comparison test were used. Correlations between continuous variables were assessed using Spearman’s rank correlation coefficient. Statistical significance was defined as follows: P ≤ 0.05 (*), P ≤ 0.01 (**), P ≤ 0.001 (***), and P ≤ 0.0001 (****). All data are presented as mean ± standard error of the mean (SEM).

## AUTHOR CONTRIBUTION

Conceptualization: V.S., D.G., P.B., D.D.

Experimental Work and Data Analysis: A.F., A.R., P.B., P.S., A.G., S.L., D.B., A.V., A.C., C.M., R.G, D.T, D.D.

Patient Recruitment and Clinical Data Analysis: A.F, A.R, B.G.-C., L.C., N.C., N.D., P. P, C. B.

Metabolic Cages Experiments: D.C., P.Z.

Circulating cfDNA Sequencing and Analysis: A.T., E.P.

Flow Cytometry Analysis: A.F, A.R, A.Z.

Vectorology: A.B

Animal Husbandry and In Vivo Sample Processing: J.I., B.R., L.M.-C.

Manuscript Writing: A.F., A.R., V.S., D.D, D.G.

Manuscript Review and Editing: All authors

Supervision: D.G., D.D., V.S.

Funding Acquisition: D.G., V.S.

## FUNDING SUPPORT

This work was supported by research grants from the Agence Nationale de la Recherche (ANR-20-CE15-0004 to V.S.), the Indo-French Centre for the Promotion of Advanced Research (IFCPAR No. 6203-1 to D.G. and V.S.), the IdEx Junior Chair Program of the University of Bordeaux (V.S.), a PhD fellowship from the Centre Hospitalier Universitaire de Bordeaux (A.F.), and the Fondation Bordeaux Université (Fonds FGLMR/AVAD pour les maladies chroniques to A.F. and V.S.), INSERM (D.C.), Nouvelle Aquitaine region (D.C.), ANR-10-EQX-008-1 optopath (D.C.). The animal platform facility of INSERM U1215 Neurocentre Magendie is supported by University of Bordeaux “investments for the future” program GPR Brain_2030.

## AKNOWLEDGMENTS

We thank the TBMCore facilities of the University of Bordeaux for their technical support in flow cytometry (Atika Zouine, Jean-Michel Griffon, and Vincent Pitard), qPCR (Xavier Gauthereau), vectorology (VECT’UB, Alice Bibeyran) and for providing adeno-associated viral vectors and John Tchen for figure design. We are also grateful to the staff of the Bordeaux A2 animal facility for their assistance in animal husbandry and experimentation. We also acknowledge the patients and the Biobank of the Aragon Health System (PT23/00146), integrated in the Platform ISCIII Biobanks and Biomodels, for their collaboration.

## CONFLICT OF INTEREST DISCLOSURE

The authors declare no conflict of interest.

## Notes

### Competing Interest Statement

The authors have declared no competing interest.

## REFERENCES

1. M. Tremmel, U.-G. Gerdtham, P. M. Nilsson, S. Saha, Economic Burden of Obesity: A Systematic Literature Review. Int J Environ Res Public Health 14, 435 (2017).

2. S. Z. Yanovski, J. A. Yanovski, Obesity. New England Journal of Medicine 346, 591–602 (2002).

3. Y. Mahamat-saleh, D. Aune, H. Freisling, S. Hardikar, R. Jaafar, S. Rinaldi, M. J. Gunter, L. Dossus, Association of metabolic obesity phenotypes with risk of overall and site-specific cancers: a systematic review and meta-analysis of cohort studies. Br J Cancer 131, 1480–1495 (2024).

4. S. M. M. Aghili, M. Ebrahimpur, B. Arjmand, Z. Shadman, M. Pejman Sani, M. Qorbani, B. Larijani, M. Payab, Obesity in COVID-19 era, implications for mechanisms, comorbidities, and prognosis: a review and meta-analysis. Int J Obes 45, 998–1016 (2021).

5. Y. S. Lee, J. Wollam, J. M. Olefsky, An Integrated View of Immunometabolism. Cell 172, 22–40 (2018).

6. S. P. Weisberg, D. McCann, M. Desai, M. Rosenbaum, R. L. Leibel, A. W. Ferrante, Obesity is associated with macrophage accumulation in adipose tissue. J Clin Invest 112, 1796–1808 (2003).

7. S. P. Weisberg, D. Hunter, R. Huber, J. Lemieux, S. Slaymaker, K. Vaddi, I. Charo, R. L. Leibel, A. W. Ferrante, CCR2 modulates inflammatory and metabolic effects of high-fat feeding. J Clin Invest 116, 115–124 (2006).

8. S. Tran, I. Baba, L. Poupel, S. Dussaud, M. Moreau, A. Gélineau, G. Marcelin, E. Magréau-Davy, M. Ouhachi, P. Lesnik, A. Boissonnas, W. Le Goff, B. E. Clausen, L. Yvan-Charvet, F. Sennlaub, T. Huby, E. L. Gautier, Impaired Kupffer Cell Self-Renewal Alters the Liver Response to Lipid Overload during Non-alcoholic Steatohepatitis. Immunity 53, 627–640.e5 (2020).

9. A. Remmerie, L. Martens, T. Thoné, A. Castoldi, R. Seurinck, B. Pavie, J. Roels, B. Vanneste, S. De Prijck, M. Vanhockerhout, M. Binte Abdul Latib, L. Devisscher, A. Hoorens, J. Bonnardel, N. Vandamme, A. Kremer, P. Borghgraef, H. Van Vlierberghe, S. Lippens, E. Pearce, Y. Saeys, C. L. Scott, Osteopontin Expression Identifies a Subset of Recruited Macrophages Distinct from Kupffer Cells in the Fatty Liver. Immunity 53, 641–657.e14 (2020).

10. K. T. Uysal, S. M. Wiesbrock, M. W. Marino, G. S. Hotamisligil, Protection from obesity-induced insulin resistance in mice lacking TNF-alpha function. Nature 389, 610–614 (1997).

11. D. Patsouris, P.-P. Li, D. Thapar, J. Chapman, J. M. Olefsky, J. G. Neels, Ablation of CD11c-positive cells normalizes insulin sensitivity in obese insulin resistant animals. Cell Metab 8, 301–309 (2008).

12. H. Wen, D. Gris, Y. Lei, S. Jha, L. Zhang, M. T.-H. Huang, W. J. Brickey, J. P.-Y. Ting, Fatty acid-induced NLRP3-PYCARD inflammasome activation interferes with insulin signaling. Nat Immunol 12, 408–415 (2011).

13. L. E. Bernstein, J. Berry, S. Kim, B. Canavan, S. K. Grinspoon, Effects of Etanercept in Patients With the Metabolic Syndrome. Arch Intern Med 166, 902–908 (2006).

14. E. J. P. van Asseldonk, R. Stienstra, T. B. Koenen, L. A. B. Joosten, M. G. Netea, C. J. Tack, Treatment with Anakinra Improves Disposition Index But Not Insulin Sensitivity in Nondiabetic Subjects with the Metabolic Syndrome: A Randomized, Double-Blind, Placebo-Controlled Study. J Clin Endocrinol Metab 96, 2119–2126 (2011).

15. D. A. Jaitin, L. Adlung, C. A. Thaiss, A. Weiner, B. Li, H. Descamps, P. Lundgren, C. Bleriot, Z. Liu, A. Deczkowska, H. Keren-Shaul, E. David, N. Zmora, S. M. Eldar, N. Lubezky, O. Shibolet, D. A. Hill, M. A. Lazar, M. Colonna, F. Ginhoux, H. Shapiro, E. Elinav, I. Amit, Lipid-Associated Macrophages Control Metabolic Homeostasis in a Trem2-Dependent Manner. Cell 178, 686–698.e14 (2019).

16. S. Daemen, A. Gainullina, G. Kalugotla, L. He, M. M. Chan, J. W. Beals, K. H. Liss, S. Klein, A. E. Feldstein, B. N. Finck, M. N. Artyomov, J. D. Schilling, Dynamic Shifts in the Composition of Resident and Recruited Macrophages Influence Tissue Remodeling in NASH. Cell Reports 34 (2021).

17. H. Shi, M. V. Kokoeva, K. Inouye, I. Tzameli, H. Yin, J. S. Flier, TLR4 links innate immunity and fatty acid–induced insulin resistance. J Clin Invest 116, 3015–3025 (2006).

18. B. Vandanmagsar, Y.-H. Youm, A. Ravussin, J. E. Galgani, K. Stadler, R. L. Mynatt, E. Ravussin, J. M. Stephens, V. D. Dixit, The NLRP3 inflammasome instigates obesity-induced inflammation and insulin resistance. Nature Medicine 17, 179–188 (2011).

19. W. L. Holland, B. T. Bikman, L.-P. Wang, G. Yuguang, K. M. Sargent, S. Bulchand, T. A. Knotts, G. Shui, D. J. Clegg, M. R. Wenk, M. J. Pagliassotti, P. E. Scherer, S. A. Summers, Lipid-induced insulin resistance mediated by the proinflammatory receptor TLR4 requires saturated fatty acid-induced ceramide biosynthesis in mice. J Clin Invest 121, 1858–1870 (2011).

20. A. Ferriere, P. Santa, A. Garreau, P. Bandopadhyay, P. Blanco, D. Ganguly, V. Sisirak, Self-Nucleic Acid Sensing: A Novel Crucial Pathway Involved in Obesity-Mediated Metaflammation and Metabolic Syndrome. Frontiers in Immunology 11 (2021).

21. H. Kwak, E. Lee, R. Karki, DNA sensors in metabolic and cardiovascular diseases: Molecular mechanisms and therapeutic prospects. Immunol Rev 329, e13382 (2025).

22. S. Nishimoto, D. Fukuda, Y. Higashikuni, K. Tanaka, Y. Hirata, C. Murata, J. Kim-Kaneyama, F. Sato, M. Bando, S. Yagi, T. Soeki, T. Hayashi, I. Imoto, H. Sakaue, M. Shimabukuro, M. Sata, Obesity-induced DNA released from adipocytes stimulates chronic adipose tissue inflammation and insulin resistance. Science Advances 2, e1501332 (2016).

23. I. Garcia-Martinez, N. Santoro, Y. Chen, R. Hoque, X. Ouyang, S. Caprio, M. J. Shlomchik, R. L. Coffman, A. Candia, W. Z. Mehal, Hepatocyte mitochondrial DNA drives nonalcoholic steatohepatitis by activation of TLR9. The Journal Of Clinical Investigation 126, 859–864 (2016).

24. X. S. Revelo, M. Ghazarian, M. H. Y. Chng, H. Luck, J. H. Kim, K. Zeng, S. Y. Shi, S. Tsai, H. Lei, J. Kenkel, C. L. Liu, S. Tangsombatvisit, H. Tsui, C. Sima, C. Xiao, L. Shen, X. Li, T. Jin, G. F. Lewis, M. Woo, P. J. Utz, M. Glogauer, E. Engleman, S. Winer, D. A. Winer, Nucleic Acid-Targeting Pathways Promote Inflammation in Obesity-Related Insulin Resistance. Cell Reports 16, 717–730 (2016).

25. A. R. Ghosh, R. Bhattacharya, S. Bhattacharya, T. Nargis, O. Rahaman, P. Duttagupta, D. Raychaudhuri, C. S. C. Liu, S. Roy, P. Ghosh, S. Khanna, T. Chaudhuri, O. Tantia, S. Haak, S. Bandyopadhyay, S. Mukhopadhyay, P. Chakrabarti, D. Ganguly, Adipose Recruitment and Activation of Plasmacytoid Dendritic Cells Fuel Metaflammation. Diabetes 65, 3440–3452 (2016).

26. X. Luo, H. Li, L. Ma, J. Zhou, X. Guo, S.-L. Woo, Y. Pei, L. R. Knight, M. Deveau, Y. Chen, X. Qian, X. Xiao, Q. Li, X. Chen, Y. Huo, K. McDaniel, H. Francis, S. Glaser, F. Meng, G. Alpini, C. Wu, Expression of STING Is Increased in Liver Tissues From Patients With NAFLD and Promotes Macrophage-Mediated Hepatic Inflammation and Fibrosis in Mice. Gastroenterology 155, 1971–1984.e4 (2018).

27. Y. Yu, Y. Liu, W. An, J. Song, Y. Zhang, X. Zhao, STING-mediated inflammation in Kupffer cells contributes to progression of nonalcoholic steatohepatitis. J Clin Invest 129, 546–555 (2019).

28. C. Li, G. Wang, P. Sivasami, R. N. Ramirez, Y. Zhang, C. Benoist, D. Mathis, Interferon-α-producing plasmacytoid dendritic cells drive the loss of adipose tissue regulatory T cells during obesity. Cell Metabolism 33, 1610–1623.e5 (2021).

29. A. D. Hildreth, E. T. Padilla, M. Gupta, Y. Y. Wong, R. Sun, A. R. Legala, T. E. O’Sullivan, Adipose cDC1s contribute to obesity-associated inflammation through STING-dependent IL-12 production. Nat Metab 5, 2237–2252 (2023).

30. P. Santa, A. Garreau, L. Serpas, A. Ferriere, P. Blanco, C. Soni, V. Sisirak, The Role of Nucleases and Nucleic Acid Editing Enzymes in the Regulation of Self-Nucleic Acid Sensing. Front Immunol 12, 629922 (2021).

31. M. Napirei, S. Wulf, D. Eulitz, H. G. Mannherz, T. Kloeckl, Comparative characterization of rat deoxyribonuclease 1 (Dnase1) and murine deoxyribonuclease 1-like 3 (Dnase1l3). Biochem J 389, 355–364 (2005).

32. A. Wilber, M. Lu, M. C. Schneider, Deoxyribonuclease I-like III is an inducible macrophage barrier to liposomal transfection. Mol Ther 6, 35–42 (2002).

33. V. Sisirak, B. Sally, V. D’Agati, W. Martinez-Ortiz, Z. B. Özçakar, J. David, A. Rashidfarrokhi, A. Yeste, C. Panea, A. S. Chida, M. Bogunovic, I. I. Ivanov, F. J. Quintana, I. Sanz, K. B. Elkon, M. Tekin, F. Yalçınkaya, T. J. Cardozo, R. M. Clancy, J. P. Buyon, B. Reizis, Digestion of Chromatin in Apoptotic Cell Microparticles Prevents Autoimmunity. Cell 166, 88–101 (2016).

34. S. M. Al-Mayouf, A. Sunker, R. Abdwani, S. A. Abrawi, F. Almurshedi, N. Alhashmi, A. Al Sonbul, W. Sewairi, A. Qari, E. Abdallah, M. Al-Owain, S. Al Motywee, H. Al-Rayes, M. Hashem, H. Khalak, L. Al-Jebali, F. S. Alkuraya, Loss-of-function variant in *DNASE1L3* causes a familial form of systemic lupus erythematosus. Nature Genetics 43, 1186–1188 (2011).

35. C. Soni, O. A. Perez, W. N. Voss, J. N. Pucella, L. Serpas, J. Mehl, K. L. Ching, J. Goike, G. Georgiou, G. C. Ippolito, V. Sisirak, B. Reizis, Plasmacytoid Dendritic Cells and Type I Interferon Promote Extrafollicular B Cell Responses to Extracellular Self-DNA. Immunity 0 (2020).

36. S. K. Tedeschi, M. Barbhaiya, S. Malspeis, B. Lu, J. A. Sparks, E. W. Karlson, W. Willett, K. H. Costenbader, Obesity and the risk of systemic lupus erythematosus among women in the Nurses’ Health Studies. Semin. Arthritis Rheum. 47, 376–383 (2017).

37. N. Hanna Kazazian, Y. Wang, A. Roussel-Queval, L. Marcadet, L. Chasson, C. Laprie, B. Desnues, J. Charaix, M. Irla, L. Alexopoulou, Lupus Autoimmunity and Metabolic Parameters Are Exacerbated Upon High Fat Diet-Induced Obesity Due to TLR7 Signaling. Front Immunol 10 (2019).

38. U. K. Dhawan, P. Bhattacharya, S. Narayanan, V. Manickam, A. Aggarwal, M. Subramanian, Hypercholesterolemia Impairs Clearance of Neutrophil Extracellular Traps and Promotes Inflammation and Atherosclerotic Plaque Progression. Arterioscler Thromb Vasc Biol 41, 2598–2615 (2021).

39. U. K. Dhawan, A. Margraf, M. Lech, M. Subramanian, Hypercholesterolemia promotes autoantibody production and a lupus-like pathology via decreased DNase-mediated clearance of DNA. Journal of Cellular and Molecular Medicine 26, 5267–5276 (2022).

40. X. Lou, Y. Hou, D. Liang, L. Peng, H. Chen, S. Ma, L. Zhang, A novel Alu-based real-time PCR method for the quantitative detection of plasma circulating cell-free DNA: Sensitivity and specificity for the diagnosis of myocardial infarction. International Journal of Molecular Medicine 35, 72–80 (2015).

41. M. D’Abbondanza, E. E. Martorelli, M. A. Ricci, S. De Vuono, E. N. Migliola, C. Godino, S. Corradetti, D. Siepi, M. T. Paganelli, N. Maugeri, G. Lupattelli, Increased plasmatic NETs by-products in patients in severe obesity. Sci Rep 9, 14678 (2019).

42. D. S. C. Han, M. Ni, R. W. Y. Chan, V. W. H. Chan, K. O. Lui, R. W. K. Chiu, Y. M. D. Lo, The Biology of Cell-free DNA Fragmentation and the Roles of DNASE1, DNASE1L3, and DFFB. The American Journal of Human Genetics 106, 202–214 (2020).

43. R. W. Y. Chan, L. Serpas, M. Ni, S. Volpi, L. T. Hiraki, L.-S. Tam, A. Rashidfarrokhi, P. C. H. Wong, L. H. P. Tam, Y. Wang, P. Jiang, A. S. H. Cheng, W. Peng, D. S. C. Han, P. P. P. Tse, P. K. Lau, W.-S. Lee, A. Magnasco, E. Buti, V. Sisirak, N. AlMutairi, K. C. A. Chan, R. W. K. Chiu, B. Reizis, Y. M. D. Lo, Plasma DNA Profile Associated with DNASE1L3 Gene Mutations: Clinical Observations, Relationships to Nuclease Substrate Preference, and In Vivo Correction. Am J Hum Genet 107, 882–894 (2020).

44. L. Serpas, R. W. Y. Chan, P. Jiang, M. Ni, K. Sun, A. Rashidfarrokhi, C. Soni, V. Sisirak, W.-S. Lee, S. H. Cheng, W. Peng, K. C. A. Chan, R. W. K. Chiu, B. Reizis, Y. M. D. Lo, Dnase1l3 deletion causes aberrations in length and end-motif frequencies in plasma DNA. Proc. Natl. Acad. Sci. U.S.A. 116, 641–649 (2019).

45. Z.-W. Li, B. Ruan, P.-J. Yang, J.-J. Liu, P. Song, J.-L. Duan, L. Wang, Oit3, a promising hallmark gene for targeting liver sinusoidal endothelial cells. Sig Transduct Target Ther 8, 1–10 (2023).

46. A. R. Saltiel, J. M. Olefsky, Inflammatory mechanisms linking obesity and metabolic disease. The Journal Of Clinical Investigation 127, 1–4 (2017).

47. J. Hartl, L. Serpas, Y. Wang, A. Rashidfarrokhi, O. A. Perez, B. Sally, V. Sisirak, C. Soni, A. Khodadadi-Jamayran, A. Tsirigos, I. Caiello, C. Bracaglia, S. Volpi, G. M. Ghiggeri, A. S. Chida, I. Sanz, M. Y. Kim, H. M. Belmont, G. J. Silverman, R. M. Clancy, P. M. Izmirly, J. P. Buyon, B. Reizis, Autoantibody-mediated impairment of DNASE1L3 activity in sporadic systemic lupus erythematosus. Journal of Experimental Medicine 218, e20201138 (2021).

48. E. Gomez-Bañuelos, Y. Yu, J. Li, K. S. Cashman, M. Paz, M. I. Trejo-Zambrano, R. Bugrovsky, Y. Wang, A. S. Chida, C. A. Sherman-Baust, D. P. Ferris, D. W. Goldman, E. Darrah, M. Petri, I. Sanz, F. Andrade, Affinity maturation generates pathogenic antibodies with dual reactivity to DNase1L3 and dsDNA in systemic lupus erythematosus. Nat Commun 14, 1388 (2023).

49. M. Jiménez-Alcázar, C. Rangaswamy, R. Panda, J. Bitterling, Y. J. Simsek, A. T. Long, R. Bilyy, V. Krenn, C. Renné, T. Renné, S. Kluge, U. Panzer, R. Mizuta, H. G. Mannherz, D. Kitamura, M. Herrmann, M. Napirei, T. A. Fuchs, Host DNases prevent vascular occlusion by neutrophil extracellular traps. Science 358, 1202–1206 (2017).

50. I. Casimiro, N. D. Stull, S. A. Tersey, R. G. Mirmira, Phenotypic sexual dimorphism in response to dietary fat manipulation in C57BL/6J mice. Journal of Diabetes and its Complications 35, 107795 (2021).

51. P. V. C. Zovico, V. H. G. Neto, F. A. Venâncio, G. P. S. Miguel, R. G. Pedrosa, F. K. Haraguchi, V. G. Baruna, Cell-Free DNA as an Obesity Biomarker. Physiol Res 69, 515–520 (2020).

52. K. Esposito, M. Ciotola, B. Schisano, R. Gualdiero, L. Sardelli, L. Misso, G. Giannetti, D. Giugliano, Endothelial microparticles correlate with endothelial dysfunction in obese women. J Clin Endocrinol Metab 91, 3676–3679 (2006).

53. A. Stepanian, L. Bourguignat, S. Hennou, M. Coupaye, D. Hajage, L. Salomon, M.-C. Alessi, S. Msika, D. de Prost, Microparticle increase in severe obesity: Not related to metabolic syndrome and unchanged after massive weight loss. Obesity 21, 2236–2243 (2013).

54. D. F. Freitas, D. F. Colón, R. L. Silva, E. M. Santos, V. H. D. Guimarães, G. H. M. Ribeiro, A. M. B. de Paula, A. L. S. Guimarães, S. T. Dos Reis, F. Q. Cunha, M. M. Antunes, G. B. Menezes, S. H. S. Santos, Neutrophil extracellular traps (NETs) modulate inflammatory profile in obese humans and mice: adipose tissue role on NETs levels. Mol Biol Rep 49, 3225–3236 (2022).

55. D. J. van der Windt, V. Sud, H. Zhang, P. R. Varley, J. Goswami, H. O. Yazdani, S. Tohme, P. Loughran, R. M. O’Doherty, M. I. Minervini, H. Huang, R. L. Simmons, A. Tsung, Neutrophil extracellular traps promote inflammation and development of hepatocellular carcinoma in nonalcoholic steatohepatitis. Hepatology 68, 1347 (2018).

56. D. S. C. Han, Y. M. D. Lo, The Nexus of cfDNA and Nuclease Biology. Trends in Genetics 37, 758–770 (2021).

57. T. H. T. Cheng, K. O. Lui, X. L. Peng, S. H. Cheng, P. Jiang, K. C. A. Chan, R. W. K. Chiu, Y. M. D. Lo, DNase1 Does Not Appear to Play a Major Role in the Fragmentation of Plasma DNA in a Knockout Mouse Model. Clinical Chemistry 64, 406–408 (2018).

58. L. Montorsi, M. J. Pitcher, Y. Zhao, C. Dionisi, A. Demonti, T. J. Tull, P. Dhami, R. J. Ellis, C. Bishop, J. D. Sanderson, S. Jain, D. D’Cruz, D. L. Gibbons, T. H. Winkler, M. Bemark, F. D. Ciccarelli, J. Spencer, Double-negative B cells and DNASE1L3 colocalise with microbiota in gut-associated lymphoid tissue. Nat Commun 15, 4051 (2024).

59. J. Spencer, S. Jain, Could tolerance to DNA be broken in the gut in systemic lupus erythematosus? Immunology Letters 270, 106937 (2024).

60. M. Takahashi, S. Ikemoto, O. Ezaki, Effect of the fat/carbohydrate ratio in the diet on obesity and oral glucose tolerance in C57BL/6J mice. J Nutr Sci Vitaminol (Tokyo) 45, 583–593 (1999).

61. J. R. Speakman, Use of high-fat diets to study rodent obesity as a model of human obesity. Int J Obes 43, 1491–1492 (2019).

62. M. Møhlenberg, E. Terczynska-Dyla, K. L. Thomsen, J. George, M. Eslam, H. Grønbæk, R. Hartmann, The role of IFN in the development of NAFLD and NASH. Cytokine 124, 154519 (2019).

63. M. Ghazarian, X. S. Revelo, M. K. Nøhr, H. Luck, K. Zeng, H. Lei, S. Tsai, S. A. Schroer, Y. J. Park, M. H. Y. Chng, L. Shen, J. A. D’Angelo, P. Horton, W. C. Chapman, D. Brockmeier, M. Woo, E. G. Engleman, O. Adeyi, N. Hirano, T. Jin, A. J. Gehring, S. Winer, D. A. Winer, Type I Interferon Responses Drive Intrahepatic T cells to Promote Metabolic Syndrome. Sci Immunol 2 (2017).

64. S. Wang, H. Ma, X. Li, X. Mo, H. Zhang, L. Yang, Y. Deng, Y. Yan, G. Yang, X. Liu, H. Sun, DNASE1L3 as an indicator of favorable survival in hepatocellular carcinoma patients following resection. Aging 12, 1171–1185 (2020).

65. L. Lei, J. Yu, B. Zhang, P. Liu, Z. Meng, Y. Song, J. Lan, B. Li, Q. Liu, Systematic profiling reveals hepatic immune and metabolic dysregulation in DNASE1L3-deficient mice. iScience 28, 114198 (2025).

66. A. Castellanos-Jankiewicz, O. Guzmán-Quevedo, V. S. Fénelon, P. Zizzari, C. Quarta, L. Bellocchio, A. Tailleux, J. Charton, D. Fernandois, M. Henricsson, C. Piveteau, V. Simon, C. Allard, S. Quemener, V. Guinot, N. Hennuyer, A. Perino, A. Duveau, M. Maitre, T. Leste-Lasserre, S. Clark, N. Dupuy, A. Cannich, D. Gonzales, B. Deprez, G. Mithieux, D. Dombrowicz, F. Bäckhed, V. Prevot, G. Marsicano, B. Staels, K. Schoonjans, D. Cota, Hypothalamic bile acid-TGR5 signaling protects from obesity. Cell Metabolism 33, 1483–1492.e10 (2021).

67. F. Andreata, C. Blériot, P. Di Lucia, G. De Simone, V. Fumagalli, X. Ficht, C. G. Beccaria, M. Kuka, F. Ginhoux, M. Iannacone, Isolation of mouse Kupffer cells for phenotypic and functional studies. STAR Protoc 2, 100831 (2021).

68. P. Bankhead, M. B. Loughrey, J. A. Fernández, Y. Dombrowski, D. G. McArt, P. D. Dunne, S. McQuaid, R. T. Gray, L. J. Murray, H. G. Coleman, J. A. James, M. Salto-Tellez, P. W. Hamilton, QuPath: Open source software for digital pathology image analysis. Sci Rep 7, 16878 (2017).

69. R. Achanta, A. Shaji, K. Smith, A. Lucchi, P. Fua, S. Süsstrunk, SLIC superpixels compared to state-of-the-art superpixel methods. IEEE Trans Pattern Anal Mach Intell 34, 2274–2282 (2012).

70. R. Meddeb, Z. A. A. Dache, S. Thezenas, A. Otandault, R. Tanos, B. Pastor, C. Sanchez, J. Azzi, G. Tousch, S. Azan, C. Mollevi, A. Adenis, S. E. Messaoudi, P. Blache, A. R. Thierry, Quantifying circulating cell-free DNA in humans. Sci Rep 9, 1–16 (2019).

71. H. Zhang, S. B. Zhang, W. Sun, S. Yang, M. Zhang, W. Wang, C. Liu, K. Zhang, S. Swarts, B. M. Fenton, P. Keng, D. Maguire, P. Okunieff, L. Zhang, B1 Sequence-based real-time Quantitative PCR: A sensitive method for direct measurement of mouse plasma DNA levels after gamma irradiation. Int J Radiat Oncol Biol Phys 74, 1592–1599 (2009).

72. D. Alpern, V. Gardeux, J. Russeil, B. Mangeat, A. C. A. Meireles-Filho, R. Breysse, D. Hacker, B. Deplancke, BRB-seq: ultra-affordable high-throughput transcriptomics enabled by bulk RNA barcoding and sequencing. Genome Biol 20, 71 (2019).

