## supplementary material for "Extracellular DNASE1L3 dysfunction fuels obesity-driven inflammation and metabolic syndrome"

#### SUPPLEMENTARY METHODS

##### Liver proteins isolation

A piece of liver was harvested from mice, placed in a solution containing PBS and protease inhibitors and disrupted using TissueLyser II (QUIAGEN). The lysate was centrifuged at 500 × g for 10 min and the supernatant was collected and stored at -80°C.

##### Isolation of human Stromal Vascular Fraction from VAT

Isolation of the SVF from VAT was carried out as previously described (*25*). Briefly, blood vessels were removed from VAT samples, and the tissue was digested in PBS supplemented with 0.075% collagenase I, 1% BSA, and 1% HEPES at 37°C. The digested tissue was then centrifuged and filtered to obtain the SVF.

##### *In vitro* macrophage differentiation

Macrophage differentiation was carried out as previously described (*25*). Briefly, $CD14^+$ monocytes were isolated from healthy PBMCs and differentiated to macrophages by culturing in the presence of recombinant human macrophage colony-stimulating factor (M-CSF) for 48h. Recombinant human IL-4 was then added to the macrophages to allow polarization to the M2-*like* phenotype for 48h. Following this, recombinant human IFN-α was added in indicated concentrations (10, 100, and 1,000 units/ml).

##### MPO and NE quantification

MPO and NE concentrations were determined using ELISA according to the manufacturer's instructions (Duoset R&D Systems, DY008, DY3174, and DY9167-05). Captured antibodies were diluted with Reagent Diluent (RD) from supplementary reagent kits (DY008), coated on 96-well microplates, and incubated overnight at RT. After three washes with wash buffer, microplates were blocked with RD for two hours. After wash step, 100µl of diluted 1/10 plasma samples, standards, and controls were added and incubated for one hour at RT. The standard curve ranged from 0.12 to 8 ng/ml. After wash step, detection antibodies diluted in RD were added for one hour. Following another wash step, microplates were incubated for 30 minutes at RT with streptavidin-HRP. After wash step, substrate solution (100µl per well) was added, and after a 10-minute incubation, the PHERAstar FS instrument, in conjunction with the PHERAstar control software, promptly read the Optical Density (O.D) at 450 nm for each well.

##### PrimeFlow™ RNA assay

Fresh PBMCs of OB patients and HD were used to perform the PrimeFlow™ RNA assay according to the manufacturer's instructions (Thermo Fischer Scientific). The probe specifically targeting the *DNASE1L3* was designed by Thermo Fisher Scientific. A control probe targeting the mRNA of *RPL13A* was also used to verify hybridization. Both probes were conjugated to an APC to allow detection by flow cytometry. Extracellular staining was performed according to the manufacturer's instructions using the following antibodies: anti-CD19 Pacific Blue, anti-CD4 SuperBright 600, anti-CD3 SuperBright 645, anti-CD304 BV711, anti-HLA-DR BV785, anti-CD123 FITC, anti-CD16 PE-EF610, anti-CD11c PE, anti-CD141 (BDCA3) PE-Cy7, anti-CD14 AF700, anti-CD1c (BDCA1) APC-Cy7, Zombie Amcyan for viability (***Supplementary Table 1***). All samples were acquired by BD LSRFortessa™ (BD Bioscience) and data were analyzed using FlowJo software version 10 (BD Biosciences).

**DNASEs activity measurement**

***Nuclei digestion assay***

Nuclei were isolating from EO771 cells using 0.05% Igepal® in hypotonic buffer, resuspended in DMEM and incubated for 2 hours at 37 °C with liver protein lysates in the presence of 2 mM $CaCl_2$ and $MnCl_2$. DNA was then extracted using the QIAamp DNA Blood Mini Kit (Qiagen), and nuclear DNA fragmentation was evaluated by agarose gel electrophoresis.

**Quantification of *DNASE1L3 expression***

Total RNA from mice liver injected with AAV or sorted immune cells was extracted and purified using the RNAeasy Plus Mini Kit (QUIAGEN) according to the manufacturer's instruction. RNA was reverse transcribed using the GoScript™ Reverse Transcription Kit (Promega). Real-time qPCR was performed using GoTaq® qPCR Master Mix (Promega). The qPCR was performed with a CFX384 thermocycler Bio-Rad real time PCR detection system. All procedures were performed according to the manufacturer's instructions. Fold change expression was calculated using $2^{-\Delta\Delta Ct}$ method with *18S* as a housekeeping gene. Primers for Human *DNASE1L3* FW : GGATCTGCTCCTTCAACGTC, RV : CCAAGCCGAGAGCTAATCAC and Mouse 18s FW : TGCCATCACTGCCATTAAG, RV : TGCTTTCCTCAACACCACATG.

**SUPPLEMENTARY FIGURES**

**Figure S1. Obese patient displays elevated levels of circulating cfDNA with a distinctive signature.**

(**A**) Ratio of circulating mitochondrial DNA (mtDNA) to nuclear DNA (nDNA) in platelet-free plasma from obese patients (OB) and healthy donors (HD). (**B**) Absolute counts of microparticles in the plasma from OB and HD, assessed by flow cytometry. (**C**) Proportion of cell-free DNA found in soluble ($MP^-$) and microparticle-associated ($MP^+$) fractions in the plasma from HD and OB. (**D**) Circulating levels of neutrophil elastase (NE) and myeloperoxidase (MPO), measured by ELISA, in the plasma from HD and OB. (**E**) Correlation between circulating nDNA and mtDNA levels with NE and MPO concentrations in OB and HD. Heatmap colors indicate correlation strength: blue = negative correlation, white = no correlation, red = positive correlation. r and p-values are shown for each pairwise comparison were determined by Spearman's test. (**F**) Linear regression analysis of circulating nDNA levels with LDL, triglycerides (TG), alanine aminotransferase (ALAT) and glucose levels. r and p-values determined by Spearman's test are indicated in each plot. (**G**) Enrichment of specific 5′ end-motifs in circulating DNA from HD (n=37), OB (n=11) and DNASE1L3-null individuals (n=2). Data are presented as mean ± SEM. Statistical significance was assessed using unpaired two-tailed t-tests. Significance is denoted as follows: ***p ≤ 0.001

**Figure S2. Obese patients exhibit reduced circulating DNASE activity.**
(**A**) DNASE1L3 mRNA expression profile in peripheral blood mononuclear cell (PBMC) subsets, compared to unstained controls (gray, Fluorescence Minus One (FMO)). Mean fluorescence intensity (MFI) values for both stained and unstained populations are indicated for each cell type. Data are representative from three independent experiments performed. (**B**) MFI of DNASE1L3 mRNA expression in PBMC subsets from obese patients (OB) and healthy donors (HD). (**C**) DNASE1L3 mRNA expression in indicated cell populations isolated from the visceral adipose tissue or PBMCs of OB individuals, measured by quantitative PCR. (**D**) DNASE1L3 mRNA expression in human macrophages differentiated from monocytes cultured with M-CSF and IL-4, in the presence or absence of increasing concentrations of recombinant human IFN-α (10, 100, and 1000 U/ml). (**E**) Correlation between plasma DNASE activity (percentage of degraded DNA) and indicated metabolic parameters: LDL cholesterol, triglycerides (TG), alanine aminotransferase (ALAT), insulin, HOMA-IR and glucose levels. Linear regression with r and p-values determined by Spearman's test are shown for each plot. (**F**) Percentage of degraded naked DNA following incubation with plasma from OB patients 3 to 12 months after bariatric surgery. (**G**) Plasma levels of anti-DNASE1L3 autoantibodies in obese patients 3 to 12 months after bariatric surgery. Data are presented as mean ± SEM. Statistical significance was determined using unpaired two-tailed t-tests or linear regression as appropriate. Significance is denoted as follows: ns = not significant; *p ≤ 0.05; **p ≤ 0.01; ***p ≤ 0.001.

**Figure S3. *Dnase1l3*-deficiency exacerbates weight gain in HFD fed male mice.**
(**A**) Cumulative food intake in WT and *Dnase1l3*-KO mice fed either a normal diet (ND) or 45% HFD. (**B-C**) Meal size over 24 hours in WT and *Dnase1l3*-KO mice in metabolic cages under ND (**B**) and 45% HFD (**C**). (**D**) Energy expenditure of WT and *Dnase1l3*-KO under 45% HFD mice during the last 4 hours of the dark phase, as indicated by the arrow in **H**. (**E-H**) Whole-

body metabolism measured over 48 hours in metabolic cages, including : (**E**) Oxygen consumption ($VO_2$), (**F**) Carbon dioxide production ($VCO_2$), (**G**) Respiratory exchange ratio (RER), and (**H**) Energy expenditure (EE) in WT and *Dnase1l3*-KO mice on ND or 45% HFD. (**I**) Body weight in WT and *Dnase1l3*-KO male mice fed ND or 60% HFD. (**J**) Cumulative body weight gain after 12 weeks on ND or 60% HFD. (**K-L**) Mass of epididymal adipose tissue (EAT) (**K**) and inguinal adipose tissue (IAT) (**L**) in WT and *Dnase1l3*-KO male mice after 12 weeks on ND or 60% HFD. (**M**) Quantification of adipocyte perimeter from 30 randomly selected cells per genotype and diet. (**N**) Body weight gain in WT and *Dnase1l3*-KO females fed ND or 60% HFD. (**O**) Cumulative body weight gain of WT and *Dnase1l*3-KO females after 12 weeks on ND or 60% HFD. (**P**) Mass of EAT in female WT and *Dnase1l3*-KO mice after 12 weeks on ND or 60% HFD. (**Q-R**) Fat (**Q**) and lean (**R**) mass measured by MRI in female mice after 12 weeks on ND or 60% HFD. Data are compiled from two independent experiments with the indicated number of mice. Results are presented as mean ± SEM. Statistical analyses were performed using unpaired two-tailed t-tests and two-way ANOVA. Significance is indicated as follows: ns = not significant; *p ≤ 0.05; **p ≤ 0.01; ***p ≤ 0.001; ****p ≤ 0.0001.

**Figure S4. *Dnase1l3*-deficiency exacerbates metabolic syndrome and MASLD in HFD fed mice**. (**A–B**) Glucose tolerance test (GTT) (**A**) and corresponding area under the curve (AUC) (**B**) in WT and *Dnase1l3*-KO male mice after 6 weeks on a normal diet (ND) or 60% HFD. (**C**) Fasting blood glucose levels after 6 weeks on ND or 60% HFD. (**D-E**) Insulin tolerance test (ITT) (**D**) and corresponding area under the curve (AUC) (**E**) in WT and *Dnase1l3*-KO male mice after 6 weeks on ND or 60% HFD. (**F**) Fasting blood insulin levels in WT and *Dnase1l3*-KO male mice after 6 weeks on ND or 60% HFD. (**G**) Homeostatic Model Assessment of Insulin Resistance (HOMA-IR) score in WT and *Dnase1l3* KO male mice after 6 weeks of ND or 60% HFD. (**H**) Plasma level of LDL cholesterol and total cholesterol in WT and *Dnase1l3*-KO male mice after 6 weeks on ND or 60% HFD. (**I-J**) GTT (**I**) and corresponding AUC (**J**) in WT and *Dnase1l3*-KO male mice after 12 weeks on ND or 60% HFD. (**K**) Fasting blood glucose levels after 12 weeks on ND or 60% HFD. (**L-M**) ITT (**L**) and corresponding AUC (**M**) in WT and *Dnase1l3*-KO male mice after 12 weeks on ND or 60% HFD. (**N**) Fasting blood insulin levels after 12 weeks of ND or 60% HFD. (**O**) HOMA-IR score in WT and *Dnase1l3*-KO male mice after 12 weeks of ND or 60% HFD. (**P**) Plasma levels of LDL cholesterol, total cholesterol after 12 weeks on ND or 60% HFD. (**Q–R**) Representative H&E-stained liver sections (**Q**) and semi-automated quantification of hepatic steatosis (**R**) in WT and *Dnase1l3*-KO male mice after 12 weeks of ND or 60% HFD. Scale bar = 400 µm. (**S-T**) GTT (**S**) and AUC (**T**) in female WT and *Dnase1l3*-KO mice after 12 weeks of ND or 60% HFD. (**U-V**) Fasting blood glucose (**U**) and insulin (**V**) levels in female WT and *Dnase1l3*-KO mice after 12 weeks of ND or 60% HFD. (**W**) HOMA-IR score in female WT and *Dnase1l3*-KO mice after 12 weeks on ND or 60% HFD. (**X**) Quantification of hepatic steatosis in female WT and *Dnase1l3*-KO mice after 12 weeks on ND or 60% HFD. Data are compiled from two independent experiments with the indicated number of mice. Results are presented as mean ± SEM. Statistical analyses were performed using unpaired two-tailed t-tests and two-way ANOVA. Significance is indicated as follows: ns = not significant; *p ≤ 0.05; **p ≤ 0.01; ***p ≤ 0.001; ****p ≤ 0.0001.

**Figure S5. *Dnase1l3*-deficiency exacerbate metabolic tissue inflammation in HFD (45%) fed mice.**

(**A**) Representative gating strategy of immune cells within the stromal vascular fraction of the epididymal adipose tissue (EAT). Macrophages : CD45$^+$ CD64$^+$ CD11b$^+$; M1-*like* : CD45$^+$ CD64$^+$ CD11b$^+$ CD11c$^+$ CD301b$^-$ ; M2-*like* : CD45$^+$ CD64$^+$ CD11b$^+$ CD11c$^-$ CD301b$^+$ ; DCs : CD45$^+$ CD64$^-$ CD11b$^-$ CD11c$^+$ MHCII$^+$ ; pDCs : CD45$^+$ CD64$^-$ CD11b$^-$ CD11c$^+$ CD317$^+$ ; T cells : CD45$^+$ TCR-β$^+$ ; B cells : CD45$^+$ B220$^+$ ; CD8 : CD45$^+$ TCR-β$^+$ CD8$^+$ ; CD4 : CD45$^+$ TCR-β$^+$ CD4$^+$ ; Tregs : CD45$^+$ TCR-β$^+$ CD4$^+$ FoxP3$^+$ ; Granulocytes : CD45$^+$ TCR-β$^-$ B220$^-$ Ly6G$^+$ ; Monocytes : CD45$^+$ TCR-β$^-$ B220$^-$ Ly6C$^+$. (**B**) Absolute counts of the indicated cell types in the EAT of WT and *Dnase1l3*-KO male mice fed a ND or 45% HFD for 12 weeks. (**C**) Absolute counts of macrophages in the EAT of WT and *Dnase1l3*-KO male mice fed a ND or 60% HFD for 12 weeks. (**D**) Frequency of M1-*like* macrophages (CD11c$^+$) and (**E**) M2-*like* macrophages (CD301b$^+$) in the EAT of WT and *Dnase1l3* KO male mice fed a ND or 60% HFD for 12 weeks. (**F**) Ratio of the number of M1-*like* to M2-*like* macrophages in the EAT of WT and *Dnase1l3*-KO male mice fed a ND or 60% HFD for 12 weeks. (**G**) Absolute counts of F4/80$^+$ CD11b$^+$ macrophages in the EAT of female WT and *Dnase1l3*-KO mice fed a ND or 60% HFD for 12 weeks. (**H**) Frequency of M1-*like* macrophages (CD45$^+$ F4/80$^+$ CD11b$^+$ CD11c and (**I**) M2-*like* macrophages (CD45$^+$ F4/80$^+$ CD11b$^+$ CD301b$^+$) in the EAT of female WT and *Dnase1l3*-KO mice fed a ND or 60% HFD for 12 weeks. (**J**) Ratio of the number of M1-*like* to M2-*like* macrophages in the EAT of female WT and *Dnase1l3*-KO mice fed a ND or 60% HFD for 12 weeks. Data were pooled from two independent experiments with the number of mice indicated. Results are presented as mean ± SEM. Statistical analyses were performed using unpaired two-tailed t-tests and two-way ANOVA. Significance is denoted as: ns = not significant; *p ≤ 0.05; **p ≤ 0.01.

**Figure S6. Adenoviral delivery of DNASE1L3 prevent MASLD development in HFD fed mice**.

(**A**) Expression of luciferase 90 days after AAV expressing luciferase (Luc) i.v injection at the indicate AAV titers. (**B**) Luciferase radiance as analyzed over time upon the i.v injection of AAV-Luc at the indicate viral titers. (**C-D**) Luciferase radiance (**C**) and expression of h*DNASE1L3* (**D**) in the indicated organs 60 days after AAV-Luc and AAV-hD1L3 i.v injection at a dose of $10^{11}$ gv/ml. (**E**) Activity of hepatic human DNASE1L3 in WT and *Dnase1l3*-KO mice 12 weeks upon i.v injection of AAV-Luc and AAV-hD1L3 at a dose of $10^{11}$ gv/ml as measured by nuclear digestion. (**F-L**) WT male mice were injected with an AAV expressing luciferase (Luc) or the human DNASE1L3 (hD1L3) at a dose of $10^{11}$ gv/ml and exposed to ND or HFD 45% during 12 weeks. (**F**) Percentage of degraded naked DNA after incubation with plasma from the indicated mice. (**G**) Cumulative food intake. (**H**) Mass of the epididymal adipose tissue (EAT) (**I**) Frequencies of $F4/80^+$ $CD11b^+$ macrophages in the EAT. (**J**) Frequency of M1-*like* macrophages ($CD45^+$ $F4/80^+$ $CD11b^+$ $CD11c^+$) and (**K**) M2-like macrophages ($CD45^+$ $F4/80^+$ $CD11b^+$ $CD301b^+$) in the EAT. (**L**) Ratio of the number of M1-*like* to M2-*like* macrophages in the EAT. Results are presented as mean ± SEM. Statistical analyses were performed using unpaired two-tailed t-tests or two-wat ANOVA. Significance is denoted as: ns = not significant; *p ≤ 0.05.

**Figure S6. Adenoviral delivery of DNASE1L3 prevent MASLD development in HFD fed mice**.

(**A**) Expression of luciferase 90 days after AAV expressing luciferase (Luc) i.v injection at the indicate AAV titers. (**B**) Luciferase radiance as analyzed over time upon the i.v injection of AAV-Luc at the indicate viral titers. (**C-D**) Luciferase radiance (**C**) and expression of h*DNASE1L3* (**D**) in the indicated organs 60 days after AAV-Luc and AAV-hD1L3 i.v injection at a dose of $10^{11}$ gv/ml. (**E**) Activity of hepatic human DNASE1L3 in WT and *Dnase1l3*-KO mice 12 weeks upon i.v injection of AAV-Luc and AAV-hD1L3 at a dose of $10^{11}$ gv/ml as measured by nuclear digestion. (**F-L**) WT male mice were injected with an AAV expressing luciferase (Luc) or the human DNASE1L3 (hD1L3) at a dose of $10^{11}$ gv/ml and exposed to ND or HFD 45% during 12 weeks. (**F**) Percentage of degraded naked DNA after incubation with plasma from the indicated mice. (**G**) Cumulative food intake. (**H**) Mass of the epididymal adipose tissue (EAT) (**I**) Frequencies of $F4/80^+$ $CD11b^+$ macrophages in the EAT. (**J**) Frequency of M1-*like* macrophages ($CD45^+$ $F4/80^+$ $CD11b^+$ $CD11c^+$) and (**K**) M2-like macrophages ($CD45^+$ $F4/80^+$ $CD11b^+$ $CD301b^+$) in the EAT. (**L**) Ratio of the number of M1-*like* to M2-*like* macrophages in the EAT. Results are presented as mean ± SEM. Statistical analyses were performed using unpaired two-tailed t-tests or two-wat ANOVA. Significance is denoted as: ns = not significant; *p ≤ 0.05.

**Supplementary Table 1. Key resources**

| REAGENT or RESOURCE | SOURCE | IDENTIFIER | CONCENTRATION (µg/ml) | DILUTION |
|---|---|---|---|---|

| | | | | |
|---|---|---|---|---|
| **Animals Diets** | | | | |
| Standard Diet A03 | SAFE | SAFE® DIET A03 | | |
| AIN93G AMF butter | SAFE | SAFE® U8978 v177 | | |
| 260HF | SAFE | SAFE®U8978 v19 | | |
| 246HF | SAFE | SAFE®U8955 v19 | | |
| **Antibodies** | | | | |
| Zombie Aqua dye AmCyan | BIOLEGEND | Cat#423102 | NA | 1/200 |
| ViaDye™ Red Fixable Viability Dye | CYTEK BIOSCIENCES | Cat#R7-60008 | NA | NA |
| Anti-mouse B220 BV421 (RA3-6B2) | BIOLEGEND | Cat#103239 | NA | 1/200 |
| Anti-mouse B220 PE Dazzle | BIOLEGEND | Cat#103258 | 0,5 | 1/400 |
| Anti-mouse B220 PE CF584 (RA3-6B2) | BD BIOSCIENCES | Cat#562313 | 0,5 | 1/400 |
| Anti-mouse CD11B APC Cy7 (M1/70) | FISHER SCIENTIFIC | Cat#A15390 | 0,25 | 1/400 |
| Anti-mouse CD11B Redfluor 710 (M1/70) | TONBO | Cat#35-0112-U500 | 1 | 1/200 |
| Anti-mouse CD11c BUV805 (N418) | BD BIOSCIENCES | Cat#749038 | 0,5 | 1/400 |
| Anti-mouse CD11c PE (N418) | BIOLEGEND | Cat#117308 | 0,4 | 1/500 |
| Anti-mouse CD16/32 (93) | FISHER SCIENTIFIC | Cat#15246827 | 5 | 1/100 |
| Anti-mouse CD19 BUV737 (1D3) | BD BIOSCIENCES | Cat#612781 | 0,33 | 1/600 |
| Anti-mouse CD206 BV711 (C068C2) | BIOLEGEND | Cat#141727 | 2 | 1/100 |
| Anti-mouse CD301B PECy7 (URA-1) | BIOLEGEND | Cat#146808 | 2 | 1/100 |
| Anti-mouse CD317 BV711 (BST2) (927) | BD BIOSCIENCES | Cat# 747604 | 1 | 1/200 |
| Anti-mouse CD4 BUV494 (RM4-4) | BD BIOSCIENCES | Cat#741051 | 0,5 | 1/400 |
| Anti-mouse CD4 FITC (RMA4-4) | BIOLEGEND | Cat#116004 | 2,5 | 1/200 |
| Anti-mouse CD44 BV510 (IM7) | BD BIOSCIENCES | Cat#563114 | 0,33 | 1/600 |
| Anti-mouse CD45 PerCP Cy5,5 (30-F11) | TONBO | Cat#65-0451 | 0,25 | 1/800 |
| Anti-mouse CD45 PerCP Cy5,5 (30-F11) | BD BIOSCIENCES | Cat#550994 | 0,25 | 1/800 |
| Anti-mouse CD62L BUV395 (MEL-14) | BD BIOSCIENCES | Cat#740218 | 1 | 1/200 |
| Anti-mouse CD64 BV605 (X54-5/7.1) | BIOLEGEND | Cat#139323 | 1 | 1/200 |
| Anti-mouse CD8 APC (53-6.7) | BD BIOSCIENCES | Cat#561093 | 1 | 1/200 |
| Anti-mouse CD8a BV786 (53-6.7) | BD | Cat#563332 | 0,5 | 1/400 |
| Anti-mouse ESAM PE (1G8/ESAM) | BIOLEGEND | Cat#136203 | 0,5 | 1/400 |
| Anti-mouse F4/80 BV650 (BM8) | BIOLEGEND | Cat#123149 | 1,33 | 1/150 |
| Anti-mouse F4/80 FITC (BM8) | BIOLEGEND | Cat#123108 | 5 | 1/100 |
| Anti-mouse FOXP3 PE (150D) | BIOLEGEND | Cat#320008 | NA | 1/200 |
| Anti-mouse FOXP3 PE-CY5 (FJK-16s) | FISHER SCIENTIFIC | Cat#15-5773-82 | 1 | 1/200 |
| Anti-mouse Ly6C BV570 (HK1.14) | BIOLEGEND | Cat#128029 | 1 | 1/200 |
| Anti-mouse Ly6C PB (HK1.4) | BIOLEGEND | Cat#128014 | 5 | 1/100 |
| Anti-mouse LY6G BV711 (1A8) | BIOLEGEND | Cat#127643 | 0,5 | 1/400 |
| Anti-mouse Ly6G Spark NIR 685 (1A8) | BIOLEGEND | Cat#127665 | 1,25 | 1/400 |

| | | | | |
|---|---|---|---|---|
| Anti-mouse MHCII PB (AF6-120,1) | BIOLEGEND | Cat#116422 | 1,25 | 1/400 |
| Anti-mouse Siglec H BV421 (551) | BD BIOSCIENCES | Cat#567815 | 1 | 1/200 |
| Anti-mouse TCRβ APC-eFluor 780 (H57-597) | FISHER SCIENTIFIC | Cat#47-5961-82 | 1,33 | 1/150 |
| Anti-mouse TIM4 APC (RMT4-54) | BIOLEGEND | Cat#130022 | 0,25 | 1/800 |
| Anti-human CD11c PE (CB16) | BD BIOSCIENCES | Cat#555392 | NA | 1/200 |
| Anti-human CD123 FITC (6H6) | BIOLEGEND | Cat#306014 | 1 | 1/200 |
| Anti-human CD14 AF700 (61D3) | FISHER SCIENTIFIC | Cat#56-0149-42 | 0,5 | 1/100 |
| Anti-human CD141 (BDCA3) PECy7 (JAA17) | FISHER SCIENTIFIC | Cat#25-1419-42 | 0,25 | 1/200 |
| Anti-human CD16 PE-EF610 (CB16) | FISHER SCIENTIFIC | Cat#61-0168-42 | 0,25 | 1/200 |
| Anti-human CD19 PB (HIB-19) | BIOLEGEND | Cat#302224 | 1,25 | 1/400 |
| Anti-human CD1c (BDCA1) APC-Cy7 (L161) | BIOLEGEND | Cat#331520 | 1 | 1/200 |
| Anti-human CD235a PE (REA175) | BD BIOSCIENCES | Cat#555570 | 2 | 1/100 |
| Anti-human CD3 SuperBright 645 (UCHT1) | FISHER SCIENTIFIC | Cat#64-0038-42 | 0,25 | 1/100 |
| Anti-human CD304 BV711 (12C2) | BIOLEGEND | Cat#354534 | 0,5 | 1/100 |
| Anti-human CD4 SuperBright 600 (SK3) | FISHER SCIENTIFIC | Cat#63-0047-42 | 0,06 | 1/200 |
| Anti-human CD41 APC (HIP8) | BIOLEGEND | Cat#303710 | NA | 1/100 |
| Anti-human HLA-DR BV785 (L243) | BIOLEGEND | Cat#307642 | 0,4 | 1/200 |
| Anti-human IgG-Fc HRP | OZYME | Cat#BETA80-104P | NA | 1/150000 |
| **Chemicals, Peptides, and Recombinant Proteins** | | | | |
| D-(+)-Glucose | SIGMA-ALDRICH | Cat#D7528 | 1g/kg | |
| Agarose | EUROMEDEX | Cat#D5 | NA | |
| Anti-FLAG M2 beads | SIGMA-ALDRICH | Cat#M8823 | NA | |
| BSA | SIGMA-ALDRICH | Cat#A7906 | NA | |
| CaCl2 | SIGMA-ALDRICH | Cat#449709 | 2mM | |
| Calf Thymus DNA | VWR | Cat#J64400.03 | 0,5µg/ml | |
| CD45 microbeads mouse | MILTENYI BIOTEC | Cat#130-052-301 | NA | |
| ClearSight DNA Stain | EUROMEDEX-BM | Cat#EUH40501 | NA | |
| Collagenase from Clostridium histolyticum | SIGMA-ALDRICH | Cat#C5138 | 8mg/ml | |
| DNASE1 | SIGMA-ALDRICH | Cat#10104159001 | 5UI/ml | |
| FBS | FISHER SCIENTIFIC | Cat#F1051 | NA | |
| Flag peptide | SIGMA-ALDRICH | Cat#F3290 | 0,25mg/ml | |
| Formalin solution, neutral buffered, 10% | SIGMA-ALDRICH | Cat#HT501128 | 0,1 | |
| FoxP3 Staining Buffer Set | FISHER SCIENTIFIC | Cat#00-5523-00 | NA | |
| Gibco™ Ack lysis buffer | FISHER SCIENTIFIC | Cat#A1049201 | NA | |
| Gibco™ DMEM Medium | FISHER SCIENTIFIC | Cat#41965-039 | NA | |
| Gibco™ HEPES | FISHER SCIENTIFIC | Cat#15630-056 | NA | |
| Gibco™ MEM NE AA | FISHER SCIENTIFIC | Cat#11140-035 | NA | |
| Gibco™ RPMI 1640 Medium | FISHER SCIENTIFIC | Cat#15460564 | NA | |
| Gibco™ Sodium pyruvate | FISHER SCIENTIFIC | Cat#11360-039 | NA | |
| GoScript™ Reverse Transcription | PROMEGA | Cat#A5001 | NA | |

| | | | | |
|---|---|---|---|---|
| GoTaq® Master Mix | PROMEGA | Cat# M7132 | 1X | |
| Igepal® CA-630 | SIGMA-ALDRICH | Cat#I8896 | 0,05% | |
| Insulin Asparte - NOVORAPID | NOVO NORDISK France | | 0,75UI/ | |
| Invitrogen™ EDTA UltraPure™ | FISHER SCIENTIFIC | Cat#11568896 | NA | |
| ISO-VET 100% | OSALIA | | 1000 mg/g | |
| MnCl2 | SIGMA-ALDRICH | Cat#244589 | 2mM | |
| Percoll® | SIGMA-ALDRICH | Cat#GE17-0891-01 | 80% / 40% | |
| Protease inibitor cocktail | SIGMA-ALDRICH | Cat#05892791001 | 1X | |
| Quant-iT™ PicoGreen™ | FISHER SCIENTIFIC | Cat#P7581 | NA | 1/200 |
| RNAlater® | SIGMA-ALDRICH | Cat#R0901 | NA | |
| TMB solution | FISHER SCIENTIFIC | Cat#10301494 | NA | |
| Tween®20 | SIGMA-ALDRICH | Cat#P1379 | 0,05% | |
| **Critical Commercial Assays** | | | | |
| PrimeFlow™ RNA Assay Kit | FISHER SCIENTIFIC | Cat#88-18005-204 | | |
| Ultra Sensitive Mouse Insulin ELISA Kit | CRYSTAL CHEM INC | Cat#90080 | | |
| U-PLEX Biomarker Group 1 (ms) Assays, QuickPlex | MSD | Cat#K15069M-21 | | |
| QIAamp DNA Blood Mini Kit | QUIAGEN | Cat#51104 | | |
| RNAeasy Plus Mini Kit | QUIAGEN | Cat#74134 | | |
| **Software, hardware and Algorithms** | | | | |
| BD Accuri C6 | BD BIOSCIENCES | | | |
| BDLSR Fortessa™ | BD BIOSCIENCES | | | |
| Bio-Rad TM CFX Manager software | BIO-RAD TM | | | |
| CFX384 thermocycler Bio-Rad real time PCR | BIO-RAD TM | | | |
| Cytek Aurora | CYTEK BIOSCIENCES | | | |
| FlowJo | TREESTAR | | | |
| GraphPad Prism | GRAPH PAD SOFTWARE | | | |
| ImageJ sofware | MacBiophotonics | | | |
| NDP View2Plus | HAMAMATSU PHOTONICS | | | |
| QuPath | UNIVERSITY OF EDINBURGH | | | |

#### Figure S1

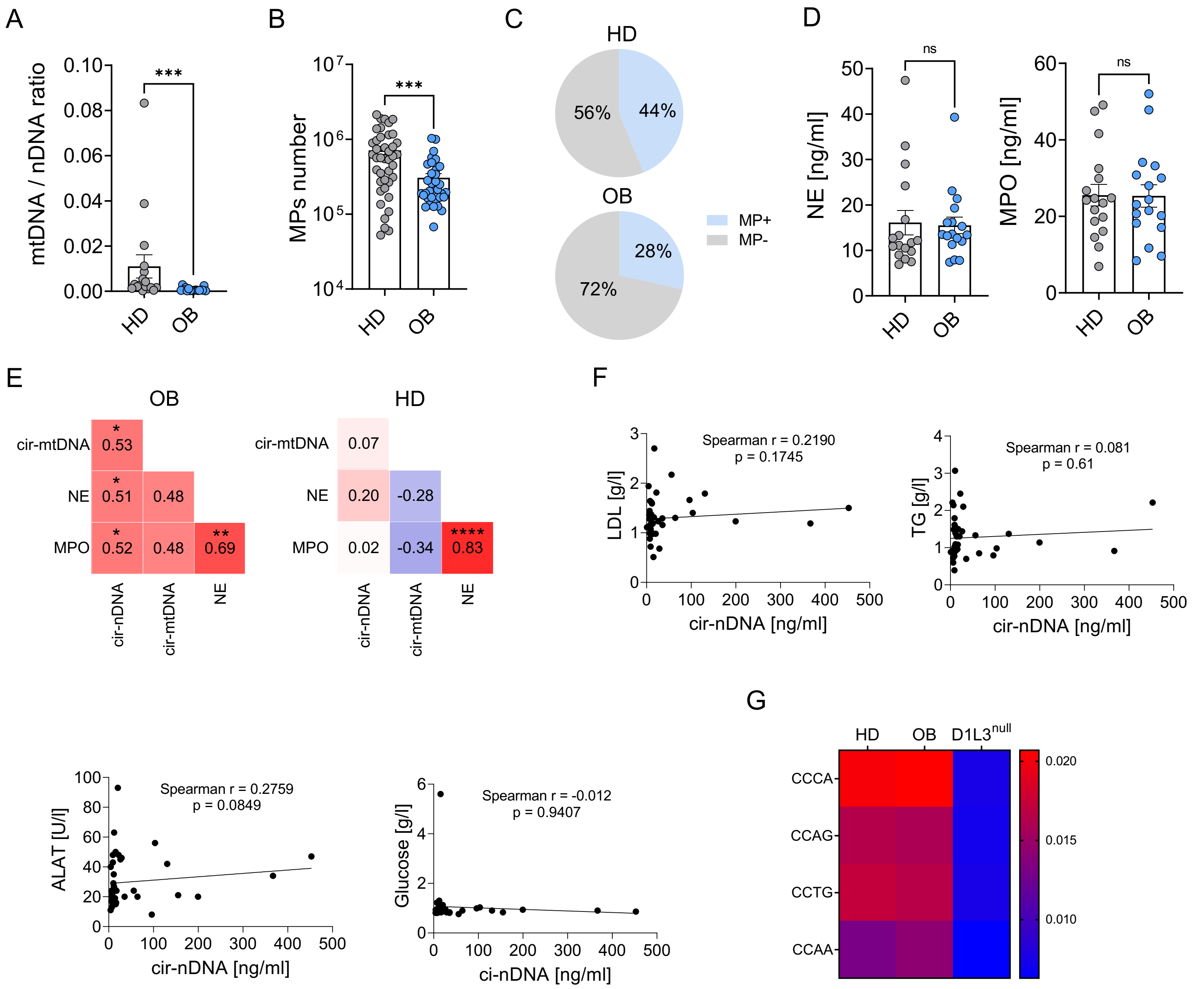


**Figure S1. Obese patient displays elevated levels of circulating cfDNA with a distinctive signature.** (**A**) Ratio of circulating mitochondrial DNA (mtDNA) to nuclear DNA (nDNA) in platelet-free plasma from obese patients (OB) and healthy donors (HD). (**B**) Absolute counts of microparticles in the plasma from OB and HD, assessed by flow cytometry. (**C**) Proportion of cell-free DNA found in soluble ($MP^-$) and microparticle-associated ($MP^+$) fractions in the plasma from HD and OB. (**D**) Circulating levels of neutrophil elastase (NE) and myeloperoxidase (MPO), measured by ELISA, in the plasma from HD and OB. (**E**) Correlation between circulating nDNA and mtDNA levels with NE and MPO concentrations in OB and HD. Heatmap colors indicate correlation strength: blue = negative correlation, white = no correlation, red = positive correlation. r and p-values are shown for each pairwise comparison were determined by Spearman's test. (**F**) Linear regression analysis of circulating nDNA levels with LDL, triglycerides (TG), alanine aminotransferase (ALAT) and glucose levels. r and p-values determined by Spearman's test are indicated in each plot. (**G**) Enrichment of specific 5′ end-motifs in circulating DNA from HD (n=37), OB (n=11) and DNASE1L3-null individuals (n=2). Data are presented as mean ± SEM. Statistical significance was assessed using unpaired two-tailed t-tests. Significance is denoted as follows: ***p ≤ 0.001

### Figure S2

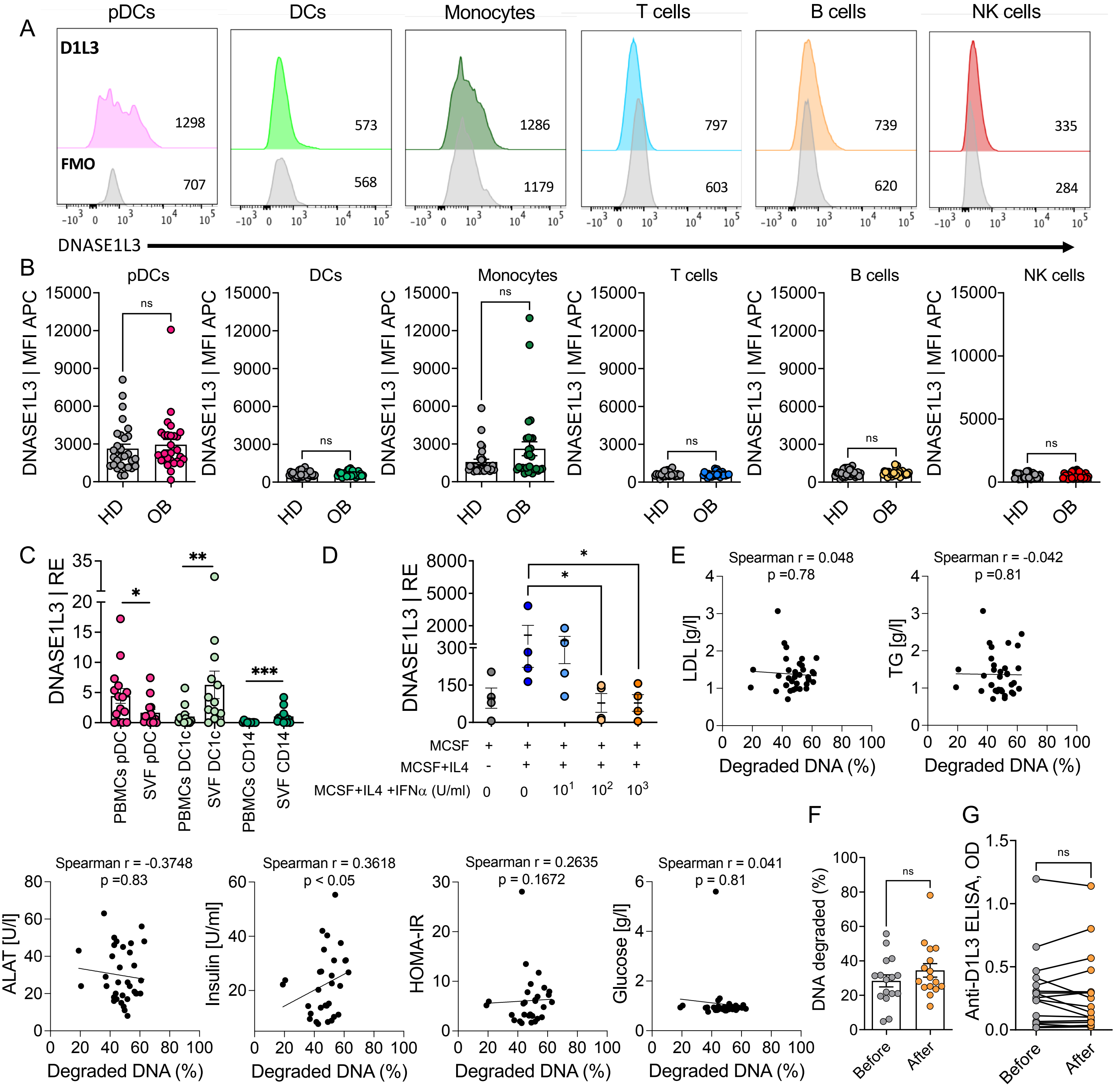


**Figure S2. Obese patients exhibit reduced circulating DNASE activity.**
(**A**) DNASE1L3 mRNA expression profile in peripheral blood mononuclear cell (PBMC) subsets, compared to unstained controls (gray, Fluorescence Minus One (FMO)). Mean fluorescence intensity (MFI) values for both stained and unstained populations are indicated for each cell type. Data are representative from three independent experiments performed. (**B**) MFI of DNASE1L3 mRNA expression in PBMC subsets from obese patients (OB) and healthy donors (HD). (**C**) DNASE1L3 mRNA expression in indicated cell populations isolated from the visceral adipose tissue or PBMCs of OB individuals, measured by quantitative PCR. (**D**) DNASE1L3 mRNA expression in human macrophages differentiated from monocytes cultured with M-CSF and IL-4, in the presence or absence of increasing concentrations of recombinant human IFN-α (10, 100, and 1000 U/ml). (**E**) Correlation between plasma DNASE activity (percentage of degraded DNA) and indicated metabolic parameters: LDL cholesterol, triglycerides (TG), alanine aminotransferase (ALAT), insulin, HOMA-IR and glucose levels. Linear regression with r and p-values determined by Spearman's test are shown for each plot. (**F**) Percentage of degraded naked DNA following incubation with plasma from OB patients 3 to 12 months after bariatric surgery. (**G**) Plasma levels of anti-DNASE1L3 autoantibodies in obese patients 3 to 12 months after bariatric surgery. Data are presented as mean ± SEM. Statistical significance was determined using unpaired two-tailed t-tests or linear regression as appropriate. Significance is denoted as follows: ns = not significant; *p ≤ 0.05; **p ≤ 0.01; ***p ≤ 0.001.

**Figure S3**

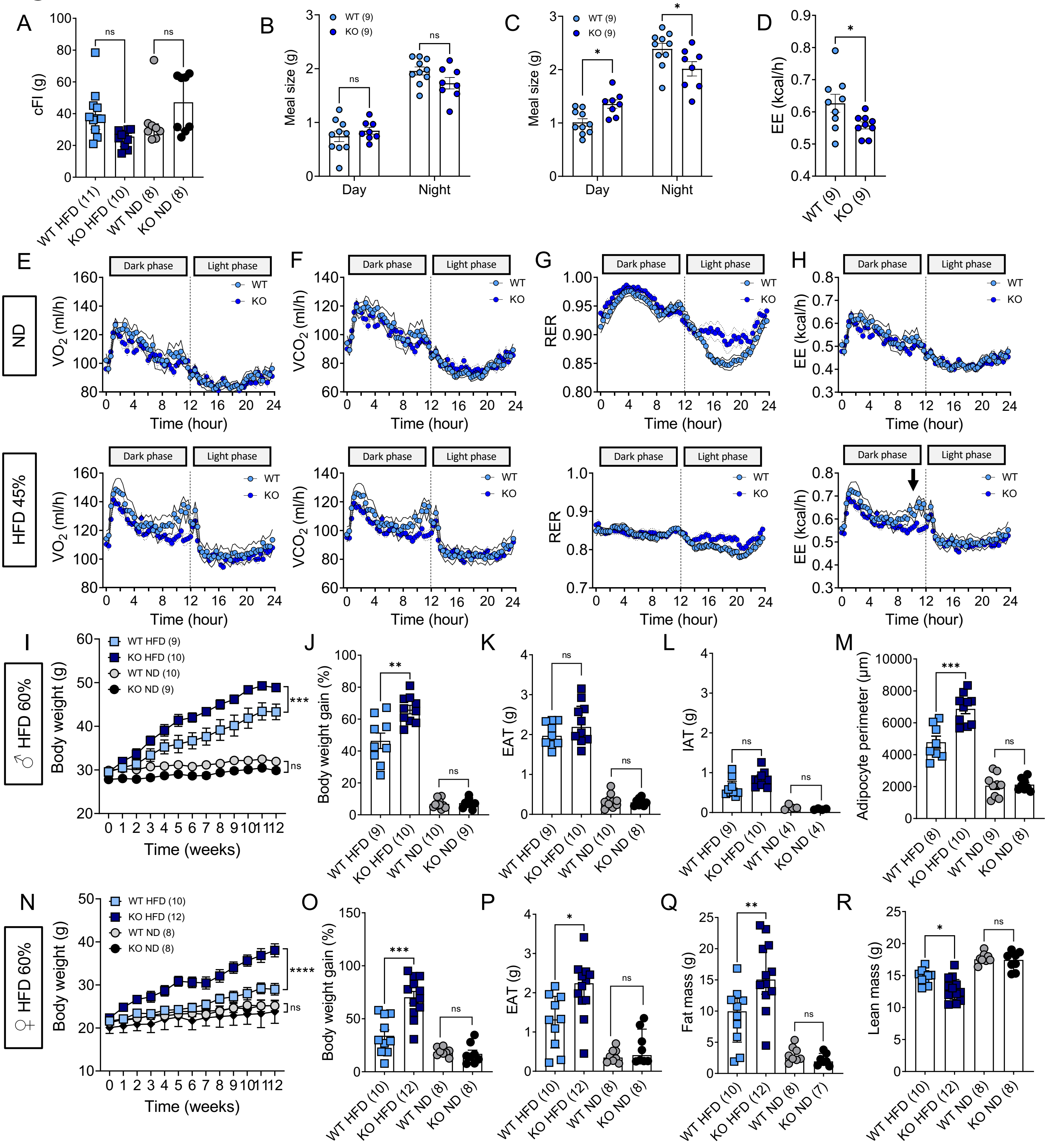


**Figure S3. *Dnase1l3*-deficiency exacerbates weight gain in HFD fed male mice.**
(**A**) Cumulative food intake in WT and *Dnase1l3*-KO mice fed either a normal diet (ND) or 45% HFD. (**B-C**) Meal size over 24 hours in WT and *Dnase1l3*-KO mice in metabolic cages under ND (**B**) and 45% HFD (**C**). (**D**) Energy expenditure of WT and *Dnase1l3*-KO under 45% HFD mice during the last 4 hours of the dark phase, as indicated by the arrow in **H**. (**E-H**) Whole-body metabolism measured over 48 hours in metabolic cages, including : (**E**) Oxygen consumption ($VO_2$), (**F**) Carbon dioxide production ($VCO_2$), (**G**) Respiratory exchange ratio (RER), and (**H**) Energy expenditure (EE) in WT and *Dnase1l3*-KO mice on ND or 45% HFD. (**I**) Body weight in WT and *Dnase1l3*-KO male mice fed ND or 60% HFD. (**J**) Cumulative body weight gain after 12 weeks on ND or 60% HFD. (**K-L**) Mass of epididymal adipose tissue (EAT) (**K**) and inguinal adipose tissue (IAT) (**L**) in WT and *Dnase1l3*-KO male mice after 12 weeks on ND or 60% HFD. (**M**) Quantification of adipocyte perimeter from 30 randomly selected cells per genotype and diet. (**N**) Body weight gain in WT and *Dnase1l3*-KO females fed ND or 60% HFD. (**O**) Cumulative body weight gain of WT and *Dnase1l3*-KO females after 12 weeks on ND or 60% HFD. (**P**) Mass of EAT in female WT and *Dnase1l3*-KO mice after 12 weeks on ND or 60% HFD. (**Q-R**) Fat (**Q**) and lean (**R**) mass measured by MRI in female mice after 12 weeks on ND or 60% HFD. Data are compiled from two independent experiments with the indicated number of mice. Results are presented as mean ± SEM. Statistical analyses were performed using unpaired two-tailed t-tests and two-way ANOVA. Significance is indicated as follows: ns = not significant; *p ≤ 0.05; **p ≤ 0.01; ***p ≤ 0.001; ****p ≤ 0.0001.

### Figure S4

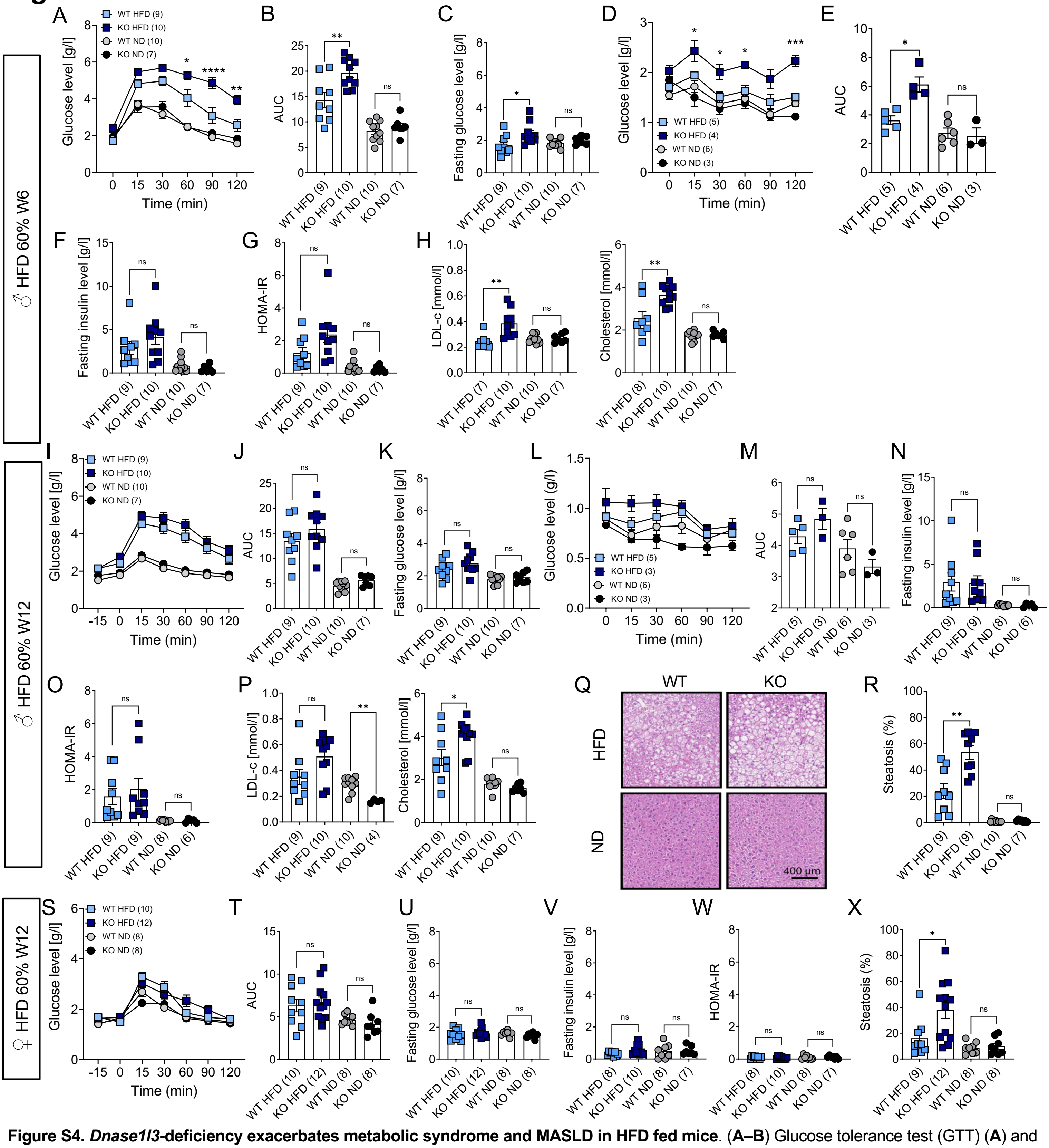


**Figure S4. *Dnase1l3*-deficiency exacerbates metabolic syndrome and MASLD in HFD fed mice**. (**A–B**) Glucose tolerance test (GTT) (**A**) and corresponding area under the curve (AUC) (**B**) in WT and *Dnase1l3*-KO male mice after 6 weeks on a normal diet (ND) or 60% HFD. (**C**) Fasting blood glucose levels after 6 weeks on ND or 60% HFD. (**D-E**) Insulin tolerance test (ITT) (**D**) and corresponding area under the curve (AUC) (**E**) in WT and *Dnase1l3*-KO male mice after 6 weeks on ND or 60% HFD. (**F**) Fasting blood insulin levels in WT and *Dnase1l3*-KO male mice after 6 weeks on ND or 60% HFD. (**G**) Homeostatic Model Assessment of Insulin Resistance (HOMA-IR) score in WT and *Dnase1l3* KO male mice after 6 weeks of ND or 60% HFD. (**H**) Plasma level of LDL cholesterol and total cholesterol in WT and *Dnase1l3*-KO male mice after 6 weeks on ND or 60% HFD. (**I-J**) GTT (**I**) and corresponding AUC (**J**) in WT and *Dnase1l3*-KO male mice after 12 weeks on ND or 60% HFD. (**K**) Fasting blood glucose levels after 12 weeks on ND or 60% HFD. (**L-M**) ITT (**L**) and corresponding AUC (**M**) in WT and *Dnase1l3*-KO male mice after 12 weeks on ND or 60% HFD. (**N**) Fasting blood insulin levels after 12 weeks of ND or 60% HFD. (**O**) HOMA-IR score in WT and *Dnase1l3*-KO male mice after 12 weeks of ND or 60% HFD. (**P**) Plasma levels of LDL cholesterol, total cholesterol after 12 weeks on ND or 60% HFD. (**Q–R**) Representative H&E-stained liver sections (**Q**) and semi-automated quantification of hepatic steatosis (**R**) in WT and *Dnase1l3*-KO male mice after 12 weeks of ND or 60% HFD. Scale bar = 400 µm. (**S-T**) GTT (**S**) and AUC (**T**) in female WT and *Dnase1l3*-KO mice after 12 weeks of ND or 60% HFD. (**U-V**) Fasting blood glucose (**U**) and insulin (**V**) levels in female WT and *Dnase1l3*-KO mice after 12 weeks of ND or 60% HFD. (**W**) HOMA-IR score in female WT and *Dnase1l3*-KO mice after 12 weeks on ND or 60% HFD. (**X**) Quantification of hepatic steatosis in female WT and *Dnase1l3*-KO mice after 12 weeks on ND or 60% HFD. Data are compiled from two independent experiments with the indicated number of mice. Results are presented as mean ± SEM. Statistical analyses were performed using unpaired two-tailed t-tests and two-way ANOVA. Significance is indicated as follows: ns = not significant; *p ≤ 0.05; **p ≤ 0.01; ***p ≤ 0.001; ****p ≤ 0.0001.

**Figure S5**

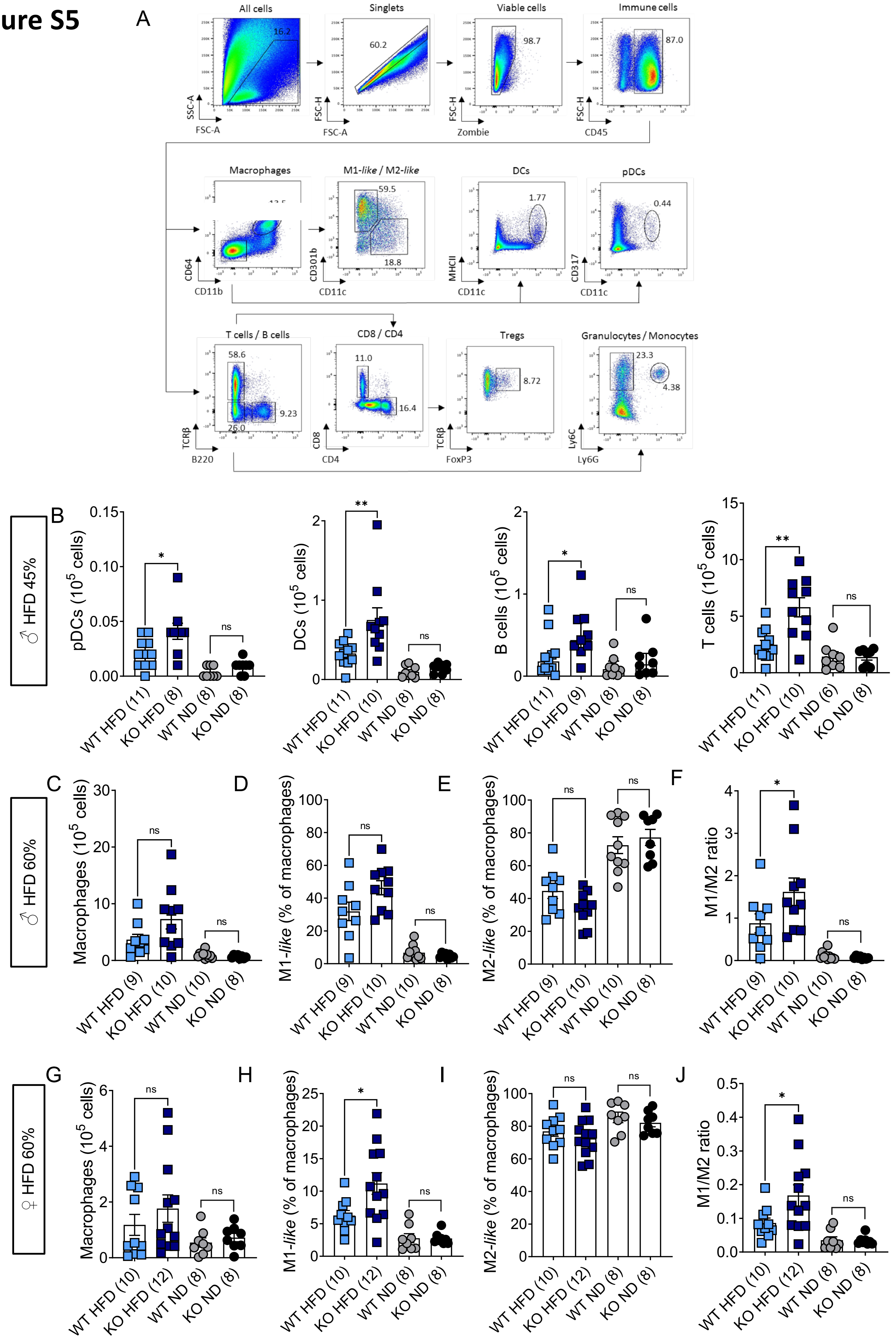

A
All cells
16.2
SSC-A
FSC-A
Singlets
60.2
FSC-H
FSC-A
Viable cells
98.7
FSC-H
Zombie
Immune cells
87.0
FSC-H
CD45
Macrophages
CD64
CD11b
M1-like / M2-like
59.5
18.8
CD301b
CD11c
DCs
1.77
MHCII
CD11c
pDCs
0.44
CD317
CD11c
T cells / B cells
58.6
9.23
26.0
TCRβ
B220
CD8 / CD4
11.0
16.4
CD8
CD4
Tregs
8.72
TCRβ
FoxP3
Granulocytes / Monocytes
23.3
4.38
Ly6C
Ly6G
B
♂ HFD 45%
pDCs (10^5 cells)
DCs (10^5 cells)
B cells (10^5 cells)
T cells (10^5 cells)
WT HFD (11)
KO HFD (8)
WT ND (8)
KO ND (8)
WT HFD (11)
KO HFD (10)
WT ND (8)
KO ND (8)
WT HFD (11)
KO HFD (9)
WT ND (8)
KO ND (8)
WT HFD (11)
KO HFD (10)
WT ND (6)
KO ND (8)
C
D
E
F
♂ HFD 60%
Macrophages (10^5 cells)
M1-like (% of macrophages)
M2-like (% of macrophages)
M1/M2 ratio
WT HFD (9)
KO HFD (10)
WT ND (10)
KO ND (8)
G
H
I
J
♀ HFD 60%
Macrophages (10^5 cells)
M1-like (% of macrophages)
M2-like (% of macrophages)
M1/M2 ratio
WT HFD (10)
KO HFD (12)
WT ND (8)
KO ND (8)

**Figure S5. *Dnase1l3*-deficiency exacerbate metabolic tissue inflammation in HFD (45%) fed mice.**

(**A**) Representative gating strategy of immune cells within the stromal vascular fraction of the epididymal adipose tissue (EAT). Macrophages : $CD45^+$ $CD64^+$ $CD11b^+$; M1-*like* : $CD45^+$ $CD64^+$ $CD11b^+$ $CD11c^+$ $CD301b^-$ ; M2-*like* : $CD45^+$ $CD64^+$ $CD11b^+$ $CD11c^-$ $CD301b^+$ ; DCs : $CD45^+$ $CD64^-$ $CD11b^-$ $CD11c^+$ $MHCII^+$ ; pDCs : $CD45^+$ $CD64^-$ $CD11b^-$ $CD11c^+$ $CD317^+$ ; T cells : $CD45^+$ TCR-$\beta^+$ ; B cells : $CD45^+$ $B220^+$ ; CD8 : $CD45^+$ TCR-$\beta^+$ $CD8^+$ ; CD4 : $CD45^+$ TCR-$\beta^+$ $CD4^+$ ; Tregs : $CD45^+$ TCR-$\beta^+$ $CD4^+$ $FoxP3^+$ ; Granulocytes : $CD45^+$ TCR-$\beta^-$ $B220^-$ $Ly6G^+$ ; Monocytes : $CD45^+$ TCR-$\beta^-$ $B220^-$ $Ly6C^+$. (**B**) Absolute counts of the indicated cell types in the EAT of WT and *Dnase1l3*-KO male mice fed a ND or 45% HFD for 12 weeks. (**C**) Absolute counts of macrophages in the EAT of WT and *Dnase1l3*-KO male mice fed a ND or 60% HFD for 12 weeks. (**D**) Frequency of M1-*like* macrophages ($CD11c^+$) and (**E**) M2-*like* macrophages ($CD301b^+$) in the EAT of WT and *Dnase1l3* KO male mice fed a ND or 60% HFD for 12 weeks. (**F**) Ratio of the number of M1-*like* to M2-*like* macrophages in the EAT of WT and *Dnase1l3*-KO male mice fed a ND or 60% HFD for 12 weeks. (**G**) Absolute counts of F4/$80^+$ $CD11b^+$ macrophages in the EAT of female WT and *Dnase1l3*-KO mice fed a ND or 60% HFD for 12 weeks. (**H**) Frequency of M1-*like* macrophages ($CD45^+$ F4/$80^+$ $CD11b^+$ CD11c and (**I**) M2-*like* macrophages ($CD45^+$ F4/$80^+$ $CD11b^+$ $CD301b^+$) in the EAT of female WT and *Dnase1l3*-KO mice fed a ND or 60% HFD for 12 weeks. (**J**) Ratio of the number of M1-*like* to M2-*like* macrophages in the EAT of female WT and *Dnase1l3*-KO mice fed a ND or 60% HFD for 12 weeks. Data were pooled from two independent experiments with the number of mice indicated. Results are presented as mean ± SEM. Statistical analyses were performed using unpaired two-tailed t-tests and two-way ANOVA. Significance is denoted as: ns = not significant; $*p \leq 0.05$; $**p \leq 0.01$.

### Figure S6

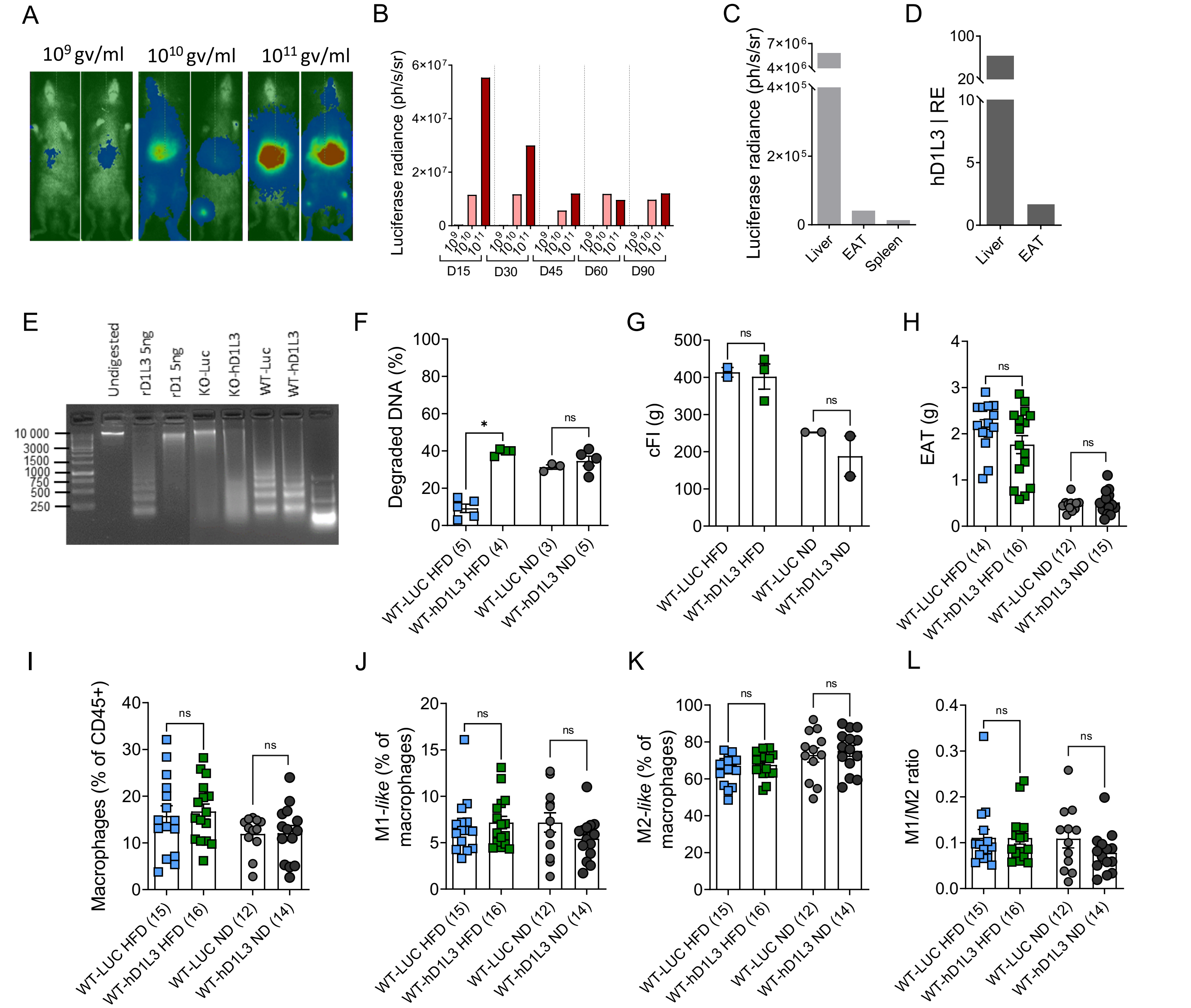


**Figure S6. Adenoviral delivery of DNASE1L3 prevent MASLD development in HFD fed mice**.
(**A**) Expression of luciferase 90 days after AAV expressing luciferase (Luc) i.v injection at the indicate AAV titers. (**B**) Luciferase radiance as analyzed over time upon the i.v injection of AAV-Luc at the indicate viral titers. (**C-D**) Luciferase radiance (**C**) and expression of h*DNASE1L3* (**D**) in the indicated organs 60 days after AAV-Luc and AAV-hD1L3 i.v injection at a dose of $10^{11}$ gv/ml. (**E**) Activity of hepatic human DNASE1L3 in WT and *Dnase1l3*-KO mice 12 weeks upon i.v injection of AAV-Luc and AAV-hD1L3 at a dose of $10^{11}$ gv/ml as measured by nuclear digestion. (**F-L**) WT male mice were injected with an AAV expressing luciferase (Luc) or the human DNASE1L3 (hD1L3) at a dose of $10^{11}$ gv/ml and exposed to ND or HFD 45% during 12 weeks. (**F**) Percentage of degraded naked DNA after incubation with plasma from the indicated mice. (**G**) Cumulative food intake. (**H**) Mass of the epididymal adipose tissue (EAT) (**I**) Frequencies of F4/80$^+$ CD11b$^+$ macrophages in the EAT. (**J**) Frequency of M1-*like* macrophages (CD45$^+$ F4/80$^+$ CD11b$^+$ CD11c$^+$) and (**K**) M2-like macrophages (CD45$^+$ F4/80$^+$ CD11b$^+$ CD301b$^+$) in the EAT. (**L**) Ratio of the number of M1-*like* to M2-*like* macrophages in the EAT. Results are presented as mean ± SEM. Statistical analyses were performed using unpaired two-tailed t-tests or two-wat ANOVA. Significance is denoted as: ns = not significant; *p ≤ 0.05.

### Figure S7

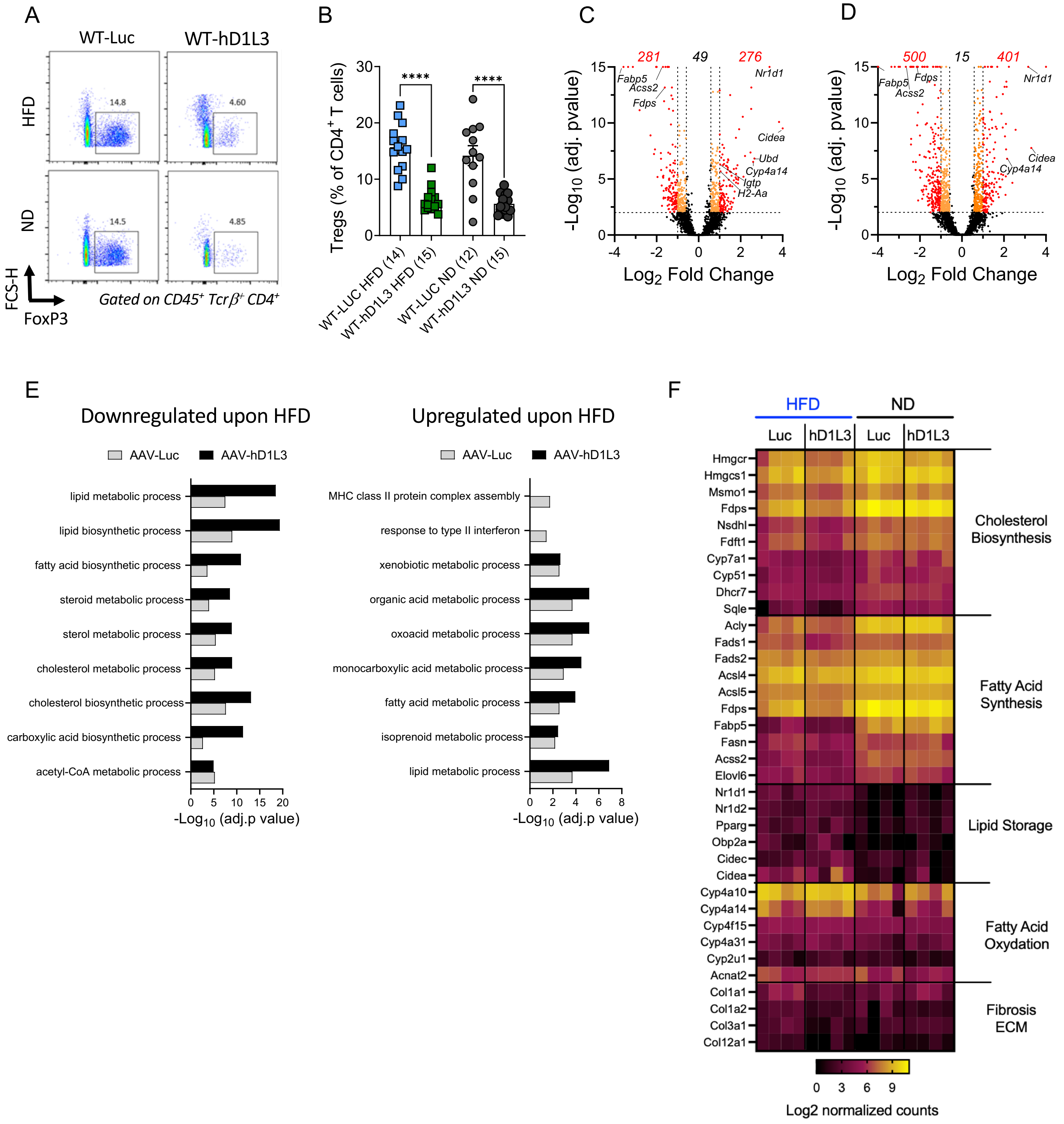


**Figure S7. Adenoviral delivery of DNASE1L3 limits liver inflammation induced by HFD.**
WT male mice were injected with an AAV expressing luciferase (Luc) or the human DNASE1L3 (hD1L3) at a dose of $10^{11}$ gv/ml and exposed to ND or HFD 45% for 12 weeks. (**A**) Representative flow cytometry plots and of regulatory T cells ($FoxP3^{+}$) in the liver after 12 weeks of the indicated diets and treatments. Numbers in plots indicate the frequencies of cells in each gate among $CD45^{+}$, $TCR\beta^{+}$, $CD4^{+}$ cells. (**B**) Frequency of regulatory T cells in the liver after 12 weeks of the indicated diets. Data were pooled from three independent experiments with the number of mice indicated. Results are presented as mean ± SEM. Statistical analyses were performed using unpaired two-tailed t-tests. Significance is denoted as: ns = not significant; $*p \le 0.05$; $**p \le 0.01$; $***p \le 0.001$; $****p \le 0.0001$. (**C-D**) Volcano plots comparing hepatic transcriptomes of WT mice fed a 45% HFD, relative to mice fed a ND that were treated with AAV-Luc (**C**) or AAV-hD1L3 (**D**). Gene ontology (GO) enrichment analysis showing HFD down- (**E**) and up-regulated (**F**) pathways in the liver of WT mice, performed separately in AAV-Luc- and AAV-hD1L3-treated mice. (**G**) Heat map showing log2 normalized expression of the indicated genes in individual mice treated with AAV-Luc or AAV-hD1L3 and fed a ND or HFD for 12 weeks.
